# Tissue damage during acute *Trypanosoma cruzi* infection is associated with reduced reparative regulatory T cell response and can be attenuated by early interleukin-33 administration

**DOI:** 10.1101/2024.02.15.580513

**Authors:** Santiago Boccardo, Constanza Rodriguez, Cintia L. Araujo Furlan, Carolina P. Abrate, Laura Almada, Camila M. S. Giménez, Manuel A. Saldivia Concepción, Peter Skewes-Cox, Srinivasa P. S. Rao, Carolina L. Montes, Adriana Gruppi, Eva V. Acosta Rodríguez

## Abstract

Tissue-repair regulatory T cells (trTregs) constitute a specialized regulatory subset renowned for orchestrating tissue homeostasis and repair. While extensively investigated in sterile injury models, their role in infection-induced tissue damage and the regulation of protective antimicrobial immunity remains largely unexplored. This investigation examines trTregs dynamics during acute *Trypanosoma cruzi* infection, a unique scenario combining extensive tissue damage with robust antiparasitic CD8+ immunity. Contrary to conventional models of sterile injury, our findings reveal a pronounced reduction of trTregs in secondary lymphoid organs and tissues during acute *T. cruzi* infection. This unexpected decline correlates with systemic as well local tissue damage, as evidenced by histological alterations and downregulation of repair-associated genes in skeletal muscle. Remarkably, a parallel decrease in systemic levels of IL-33, a crucial factor for trTregs survival and expansion, was detected. We found that early treatment with systemic recombinant IL-33 during infection induces a notable surge in trTregs, accompanied by an expansion of type 2 innate lymphoid cells and parasite-specific CD8+ cells. This intervention results in a mitigated tissue damage profile and reduced parasite burden in infected mice. These findings shed light on trTregs biology during infection-induced injury and demonstrate the feasibility of enhancing a specialized Tregs response without impairing the magnitude of effector immune mechanisms, ultimately benefiting the host. Furthermore, this study settles groundwork of relevance for potential therapeutic strategies in Chagas’ disease and other infections.

**AUTHOR SUMMARY:** Chagas’ disease, caused by the protozoan *Trypanosoma cruzi*, induces severe organ damage caused by the interplay between the parasite and the immune response. In our investigation, we delved into the role of tissue-repair regulatory T cells (trTregs) during the acute phase of *T. cruzi* infection in mice. Surprisingly, we observed a reduction in trTregs during the peak of tissue damage, contrary to their usual accumulation after injury in other contexts. This decline aligned with decreased levels of interleukin-33, a critical factor for trTregs survival. Administering interleukin-33 at early infection times not only boosted trTregs but also expanded other reparative and antiparasitic immune cells. Consequently, these treated mice exhibited reduced damage and lower parasite levels in tissues. Our findings offer insights into trTregs’ behavior during infection-induced injury, suggesting a promising avenue for therapeutic interventions in Chagas’ disease and related conditions. This study lays the groundwork for potential strategies that balance the immune response, supporting tissue repair without compromising the ability to control the infection, which could have broader implications for infectious diseases and tissue damage-related pathologies.

## INTRODUCTION

Regulatory T cells (Tregs) are CD4+ T lymphocytes with immunoregulatory properties characterized by the expression of the transcription factor Forkhead box P3 (Foxp3) [1]. The immunomodulatory role of Tregs has been extensively described across diverse biological processes, encompassing tolerance maintenance, autoimmunity and allergy, as well as cancer, infectious and immunometabolic diseases [2]. This wide spectrum of activities underscores the adaptability of Tregs in regulating various effector responses, including Th1, Th2, and Th17 immunity. This phenomenon, recognized as Tregs specialization, enables them to tailor their regulatory capacity to distinct scenarios and the specific immune profiles requiring modulation [3].

Recently, it has been identified a separate subset of Tregs that goes beyond their classic suppressive function to specialize in maintaining tissue homeostasis and promoting repair following damage [4]. These cells, referred to as tissue repair Tregs (trTregs), were initially believed to be confined to non-immune tissues, but have subsequently been found in lymphoid organs as well [5]. Tissue repair Tregs exhibit a high level of activation and share a phenotypic and transcriptomic core signature across different tissues. Nevertheless, they also possess unique characteristics and functions shaped by the microenvironment of their respective residing sites. The trTregs program is established through a stepwise process, beginning with an initial commitment in peripheral lymphoid organs and culminating in final differentiation within tissues [6–8]. Regarding function, trTregs located in adipose tissue adapt to regulate metabolism and restrain obesity [9]. This cell subset also orchestrates tissue regeneration and homeostasis in the colon [10], participates in skeletal muscle (SM) regeneration and control of fibrosis [11,12], facilitates wound healing and hair growth in the skin [13,14] and promotes myelin regeneration in the central nervous system (CNS) [15]. Throughout these contexts, the survival, expansion, and acquisition of tissue-repair properties by trTregs have been demonstrated to rely on IL-33, an alarmin belonging to the IL-1 family, released upon cellular damage [16]. Correspondingly, trTregs express the specific IL-33 receptor subunit, ST2 [5].

The majority of studies on trTregs have focused on their behavior under homeostatic conditions or in models of sterile injury, where they accumulate locally to facilitate tissue healing [17]. Nonetheless, limited information exists concerning trTregs behavior and their dependence on the IL-33/ST2 axis for controlling tissue damage induced by infections, which exposes these cells to a distinct environment. Currently, only a handful of studies have demonstrated local trTregs accumulation and their role in promoting tissue repair during acute infections caused by influenza virus [18], herpes simplex virus [19], and cytomegalovirus [20]. Intriguingly, trTregs accumulation in the lung and cornea appeared to rely more on IL-18, another IL-1 family cytokine, than on IL-33 signaling. Additionally, trTregs increased in the liver during S. japonicum helminth infection, where they significantly ameliorated tissue pathology [21]. In contrast, ST2+ Tregs that accumulated in the intestinal lamina propria of chronically HIV-infected patients had a limited role in tissue repair, as these individuals exhibited heightened epithelial permeability and tissue fibrosis [22]. In protozoan infections, different roles for trTregs have been reported, with their protective function noted in cerebral malaria [23], while their relevance in SM pathology during toxoplasmosis appeared negligible [24]. Despite these emerging reports, the role of trTregs and the IL-33/ST2 axis in tissue repair and homeostasis during infections remains poorly delineated. Few studies have examined the impact of manipulating the abundance of trTregs on antimicrobial immunity and pathogen load control. Given that trTregs are recognized for their strong suppressive potential [4,25], further exploration of their influence on effector immune responses is necessary. This exploration will contribute to a better understanding of the interplay between microbial persistence, tissue damage and repair, and the underlying immunopathology that characterizes certain chronic infections.

Chagas’ Disease (American Trypanosomiasis) is a chronic infection caused by the protozoan parasite *Trypanosoma cruzi*. It is endemic in Latin America, but cases are also reported in non-endemic regions, with approximately 6-7 million people estimated to be infected worldwide [26]. During the acute phase, the parasite invades cells from tissues such as muscle, liver, gut, lymph nodes, spleen, and the CNS, where it actively replicates, inducing cell death and tissue damage. Consequently, the acute phase of this infection is associated with high parasitemia and nonspecific symptoms. Type 1 immunity, characterized by elevated levels of proinflammatory cytokines like IFN-γ [27–29], along with innate and adaptive immune cells like NK cells, inflammatory macrophages, and CD8+ T lymphocytes [28,30,31], work together to minimize parasite replication and burden. However, this immune response is insufficient to completely eradicate the parasite from the host, leading to chronic infection. During the chronic phase, around 30% of infected individuals will develop specific symptoms related to digestive or cardiac pathology after 10-30 years [32]. Presently, evidence suggests that both parasite persistence and a sustained inflammatory environment mediate the characteristic tissue damage observed in this disease during the chronic phase [33]. In addition to the classical pathology, muscular pain and weakness are frequently observed in both acute and chronic Chagas’ patients [34–36]. Furthermore, an association between SM parasitism and myositis, with structural alterations of muscular fibers, has been demonstrated in chronically infected humans [37,38], as well as in mouse models of acute and chronic infection [39–43].

In this study, we used a mouse model of acute *T. cruzi* infection to investigate trTregs and their association with tissue damage and protective immunity. We examined trTregs dynamics in this infection scenario, characterized by a unique combination of elevated systemic tissue injury and a limited Tregs response [44]. We found reduced trTregs numbers in target tissues that correlated with reduced systemic IL-33 levels. By supplementation with recombinant IL-33, we dissected the impact of trTregs and other IL-33-responsive immune subsets on tissue damage, parasite control, and infection progression. Through this work, we shed light on the intricate balance between microbicidal and regenerative responses driven by trTregs and the IL-33/ST2 axis, an area still inadequately defined in infections and of particular relevance to the progression of acute Chagas’ Disease.

## RESULTS

### Systemic tissue damage during acute *T. cruzi* infection associates with tissue parasitism and immune infiltrate

As a first step to dissect the ability of trTregs to control damage, we initially assessed the extent and progression of tissue injury in our experimental model of acute *T. cruzi* infection. As established in our lab [45], intraperitoneal injection of 5,000 trypomastigotes (Tulahuen strain) induces a peak of parasitemia at 21 days post-infection (dpi) (Fig 1A). This is accompanied by the maximum parasite load in tissues such as SM, heart, spleen, and liver (Fig 1B), as well as the peak of immune cell expansion in the spleen (Fig 1C). In agreement with previous findings [39,46], histological examination of SM at the infection peak revealed the presence of parasite nests and diffuse mononuclear infiltrate associated with necrosis and calcification of muscular fibers, absent in samples from non-infected (NI) mice (Fig 1D). Quantification of the immune infiltrate in SM showed that leucocyte counts were also at their maximum at 21 dpi (Fig 1E).

**Fig 1:**
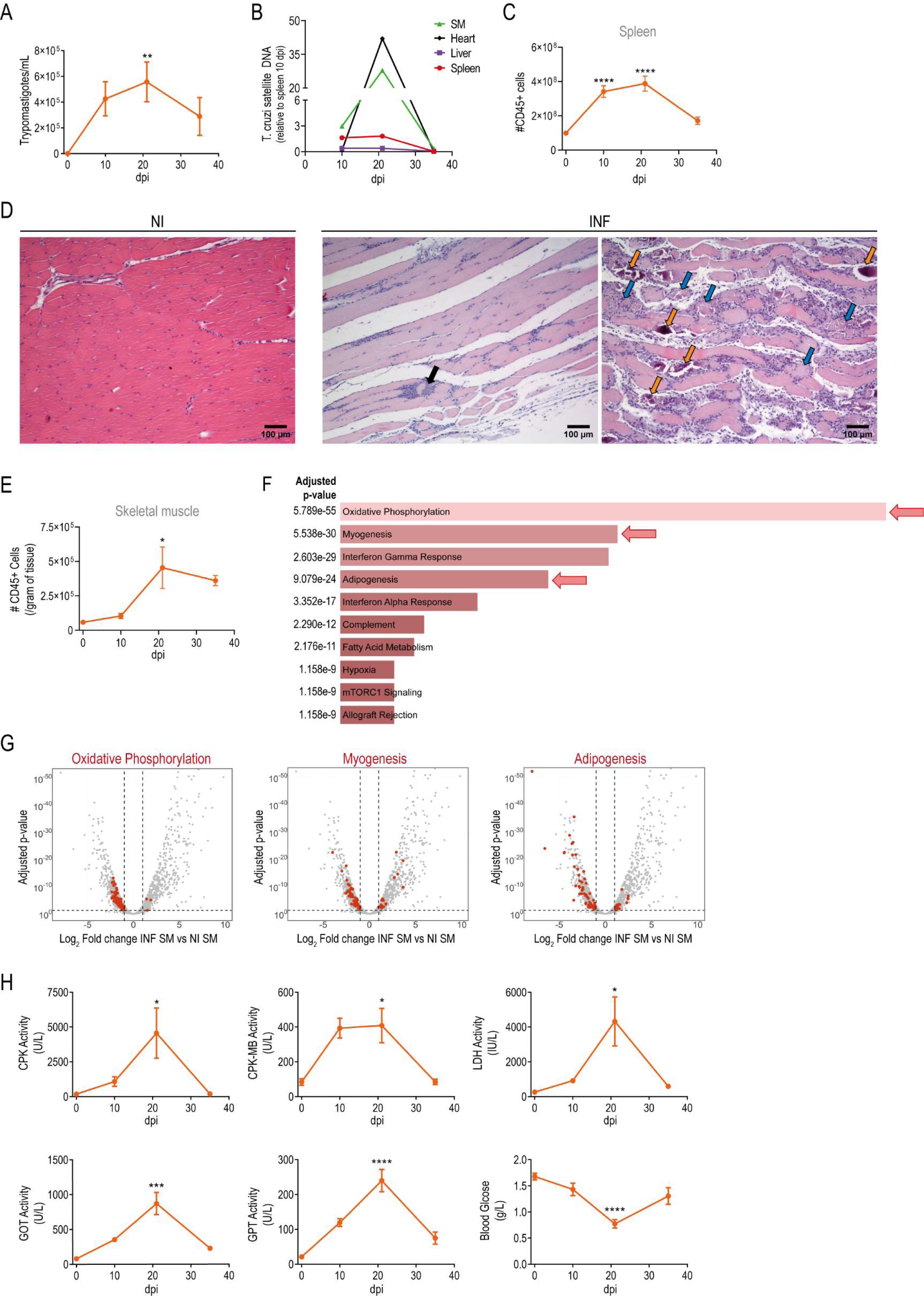
Characterization of muscular and systemic tissue damage during acute *T. cruzi* infection. Foxp3-GFP mice were infected with *T. cruzi* and tissue damage and infection progression were evaluated at different days post infection (dpi). (A) Kinetics of parasite counts in blood. (B) Kinetics of *T. cruzi* satellite DNA quantification in skeletal muscle (SM), heart, liver and spleen. (C) Kinetics of total spleen CD45+ cell count. (D) Representative Hematoxylin-Eosin stain of quadriceps muscle from non-infected (NI) and infected (INF) (21 dpi) mice (N = 4-7). Black arrow: parasites nest, blue arrows: necrotic muscle fibers, orange arrows: calcified muscle fibers. Magnification = 10X. INF images represent different areas from the same sample. (E) Kinetics of total SM CD45+ cell count. (F-G) Whole quadriceps SM RNAseq data analysis from NI and INF animals; N = 3 per group. (F) Non-supervised pathway analysis of the differentially expressed genes between INF and NI SM. Bars show the top-ten pathways upregulated in INF SM with red arrows highlighting pathways related to SM physiology. (G) Volcano plots display differentially expressed genes between INF and NI SM. According to (F), genes associated with oxidative phosphorylation (left), myogenesis (center) and adipogenesis (right) pathways are highlighted in red. (H) Kinetics of plasma CPK, CPK-MB, LDH, GOT and GPT activities, and glucose concentration. (A, C, E and H) Data is presented as mean ± SEM; N = 4-15 per dpi. Statistical significance was determined by one-way ANOVA. P values are relative to 0 dpi: *p < 0.05; **p < 0.01; ***p < 0.001; ****p < 0.0001. (B) Data are presented as mean and values are normalized to spleen tissue parasitism at 10 dpi; N = 4-5 per dpi. (A-E and H) Data were collected from 2-3 independent experiments.

To further elucidate the impact of *T. cruzi* infection on muscle physiology, we conducted a whole-tissue RNAseq comparing infected (INF) versus NI quadriceps. The transcriptome analysis identified 1621 differentially expressed genes (DEGs) between both conditions. Non-supervised pathway analysis of the DEGs using EnrichR revealed that, in addition to pathways associated with immune responses such as interferon-gamma, interferon-alpha and complement responses, several pathways related to muscle physiology such as oxidative phosphorylation, myogenesis and adipogenesis were among the most significantly enriched pathways (Fig 1F). As expected, volcano plots revealed that most genes associated with the pathways linked to immune responses were upregulated by the infection (S1A Fig and S1 Table). In contrast, the majority of genes linked to muscle physiology pathways were downregulated (Fig 1G and S1 Table), supporting the notion that acute infection disrupted SM homeostasis. Consistent with histological and transcriptomic evidences of SM damage, we found increased plasma activity of creatine phosphokinase (CPK) and creatine phosphokinase of muscle and brain (CPK-MB) at 21 dpi compared to NI mice (Fig 1H). The alteration of additional markers of systemic damage such as increased activity of lactate dehydrogenase (LDH), glutamic oxaloacetic transaminase (GOT) and glutamic pyruvic transaminase (GPT), along with hypoglycemia, indicated affection of various target tissues beyond SM, as previously reported by our group [44,47]. As expected, the greatest tissue alteration coincides with highest parasitemia counts and tissue parasitism (Figs 1A and 1B) as well as with the peak of immune cells expansion in the spleen (Fig 1C) and maximum immune infiltration in SM (Fig 1E), and other target tissues such as heart and liver (S1B Fig). Given these features, the acute phase of *T. cruzi* infection emerged as an instrumental setting to study trTregs roles during the infection.

### *Bona fide* trTregs are reduced in target tissues and lymphoid organs at the peak of infection

Given that the greatest alteration of biochemical markers of tissue damage was observed at the peak of infection, we selected 21 dpi as the initial time point to evaluate by flow cytometry the presence of total Tregs as well as of trTregs in different *T. cruzi* target sites. As previously documented [44], *T. cruzi* infected mice exhibited reduced frequencies of total Tregs in spleen and liver around the peak of the infection (S2A and S2B Figs). An even more pronounced decrease was observed when analyzing other peripheral tissues known to be common parasite targets, such as SM and the heart. We then quantified trTregs that were identified within the Foxp3+ Tregs population according to their co-expression of ST2 and KLRG-1, as proposed by Delacher et al. [5]. We detected a notable reduction in the presence of trTregs, both in terms of frequency within total Tregs and in absolute numbers, in SM, liver and spleen from INF animals compared to NI controls (Figs 2A and 2B). This subset was not evaluated in the heart due to the remarkably low infiltration of total Tregs, which impeded a more in-depth examination of this organ.

**Fig 2:**
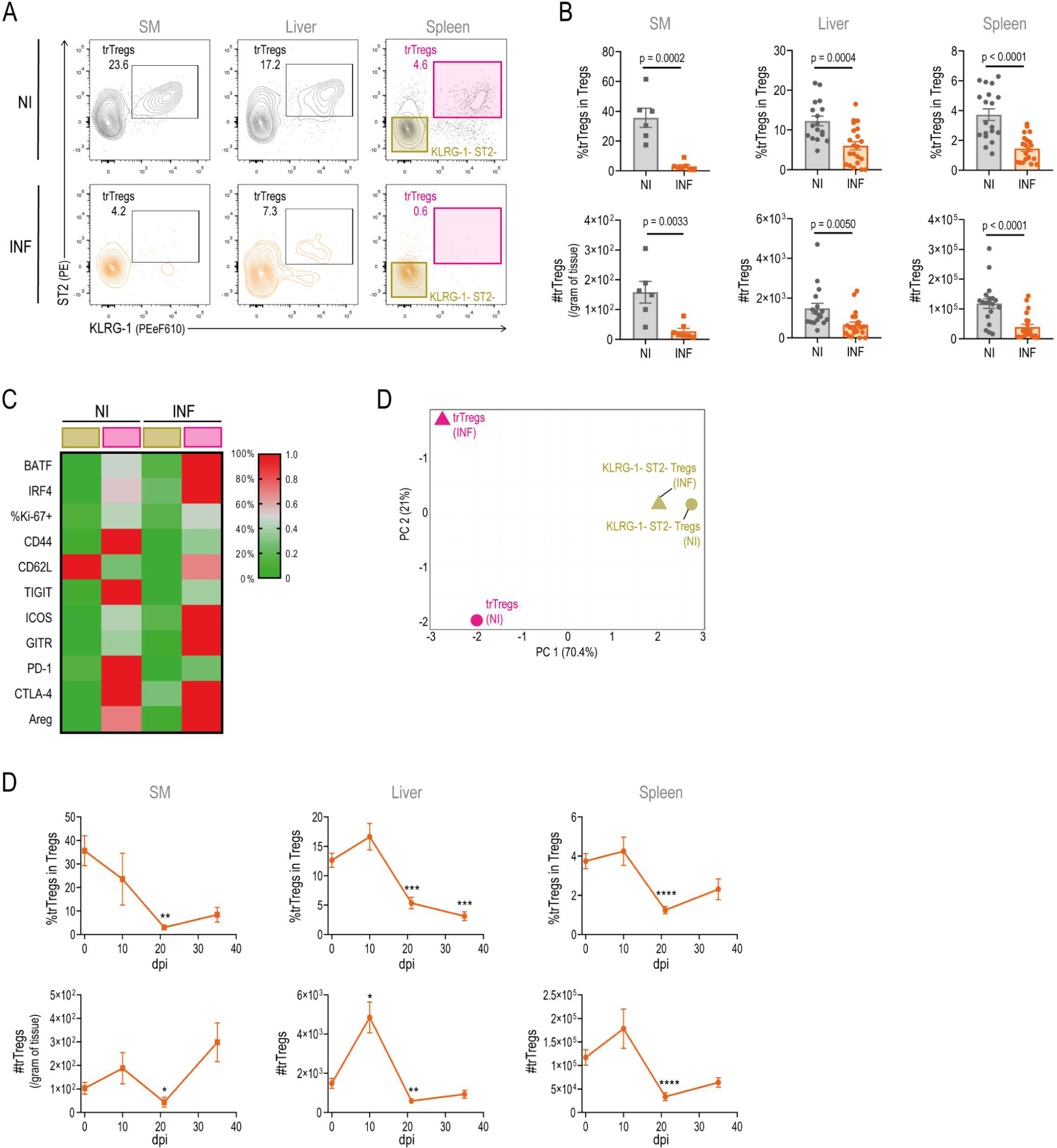
Tissue repair Tregs are reduced in target tissues and lymphoid organs during acute *T. cruzi* infection. Tissue repair Tregs (trTregs) were studied by flow cytometry in tissues from *T. cruzi* infected Foxp3-GFP mice at different days post infection (dpi). (A) Representative dot plots showing ST2 and KLRG-1 staining in total Tregs present in skeletal muscle (SM), liver and spleen obtained from non-infected (NI) and infected (INF) (21 dpi) mice. ST2+ KLRG-1+ cells (pink gate) were defined as trTregs. (B) Graphs displaying trTregs frequency within total Tregs (upper row) and absolute number (bottom row) in different tissues from NI and INF animals. Bars indicate the mean ± SEM. For SM, squares represent pools with N=3 mice. For spleen and liver, circles represent individual mice. (C) Heatmap displaying the relative expression or frequency of the indicated markers in splenic trTregs (pink gate) or ST2-KLRG-1-Tregs (golden gate) as defined in (A), evaluated in NI and INF animals (N=5-6). (D) Principal component analysis of data presented in (C). (E) Kinetics of trTregs frequency within total Tregs (upper row) and absolute number (bottom row) in SM, liver and spleen. Data are presented as mean ± SEM; N = 6-21 per dpi. (B, E) For SM, cell counts are normalized to tissue weight. Statistical significance was determined by Unpaired t test (B) and Kruskal-Wallis test (E). P values in (B) represent pairwise comparisons, while in (E) are relative to 0 dpi: *p < 0.05; **p < 0.01; ***p < 0.001; ****p < 0.0001. (A-E) Data were collected from 2-3 independent experiments.

Afterwards, we determined whether trTregs identified through the co-expression of ST2 and KLRG-1 in the context of the infection possessed distinctive phenotype of *bona fide* tissue repair cells, setting them apart from classic ST2-KLRG-1-lymphoid-like Tregs as previously described in other experimental settings [5,7,48–52]. Thus, we compared the expression of transcription factors (BATF, IRF4and Ki-67) and surface molecules (CD44, CD62L, TIGIT, ICOS, GITR, PD-1 and CTLA-4), as well as amphiregulin (Areg) production by flow cytometry in trTregs and ST2-KLRG-1-Tregs from spleen of INF and NI mice identified as depicted in Fig 2A. As shown in the representative histograms from S3 Fig and summarized in the heatmap in Fig 2C, the expression levels of most evaluated markers, with the exception of CD62L, were higher in trTregs compared to ST2-KLRG-1-Tregs obtained from the spleen of NI mice. Notably, trTregs from the spleen of INF mice also exhibited a *bona fide* trTregs phenotype, displaying even stronger phenotypic features in comparison to their ST2-KLRG-1-Tregs counterparts, with the highest expression of certain markers as BATF, IRF4, ICOS and GITR along with reduced CD44, TIGIT and PD1 levels. A principal component analysis (PCA) of the phenotypic data demonstrated that while ST2-KLRG-1-Tregs from spleens of NI and INF mice clustered closely, trTregs segregated apart along PC1, that explains around 70% of the variance among the samples (Fig 2D). Furthermore, trTregs from NI and INF mice showed some differences between them at expense of PC2, which explains around 20% of the variance mainly driven by CD62L expression.

As trTregs have been shown to increase once the tissue damage is well established [11,12,18–21], we evaluated trTregs response along the acute phase of the infection (Fig 2E). Despite observing a transient increase in trTregs numbers in the liver at 10 dpi, we found a significant reduction in the frequency and absolute number of this cell subset at 21 dpi in all evaluated tissues. It is noteworthy that while trTregs absolute numbers recover at later time points, they do not substantially increase as observed in other injury models.

In summary, our results indicate that during acute *T. cruzi* infection, despite the severe systemic tissue damage, trTregs are particularly reduced within an already restricted total Treg pool. The few remaining trTregs found at the peak of infection share several characteristics with *bona fide* trTregs, presenting particularities likely as a consequence of the infection.

### Systemic IL-33 levels are reduced during acute *T. cruzi* infection

It is established that trTregs development is a multi-step process that initiates with a specialization commitment in secondary lymphoid organs and culminates after migration into residence tissues [6–8]. Remarkably, IL-33 has been shown to play a role in all these different stages [53]. In light of the general decline in trTregs numbers despite the elevated tissue damage around the peak of acute infection, we performed a kinetic study to assess IL-33 concentrations at both systemic and peripheral sites. Our analysis revealed that plasma IL-33 concentration was detectable in NI mice and diminished during the course of acute *T. cruzi* infection, showing a statistically significant decrease at 21 dpi, followed by a return to baseline levels at 35 dpi (Fig 3A). A similar trend was observed when quantifying IL-33 in the spleen homogenates (S4A Fig). In contrast, the quantification of total IL-33 in homogenates from target tissues showed an increased concentration at 21 and 35 dpi in SM (Fig 3A) while it remained relatively constant throughout the tested period in the liver (S4A Fig). These results indicate that, despite a conservation or increase in IL-33 levels in peripheral non-lymphoid tissues, its concentration is reduced at systemic level and in secondary lymphoid tissues, suggesting that the initial steps of trTregs development may be affected during *T. cruzi* infection.

**Fig 3:**
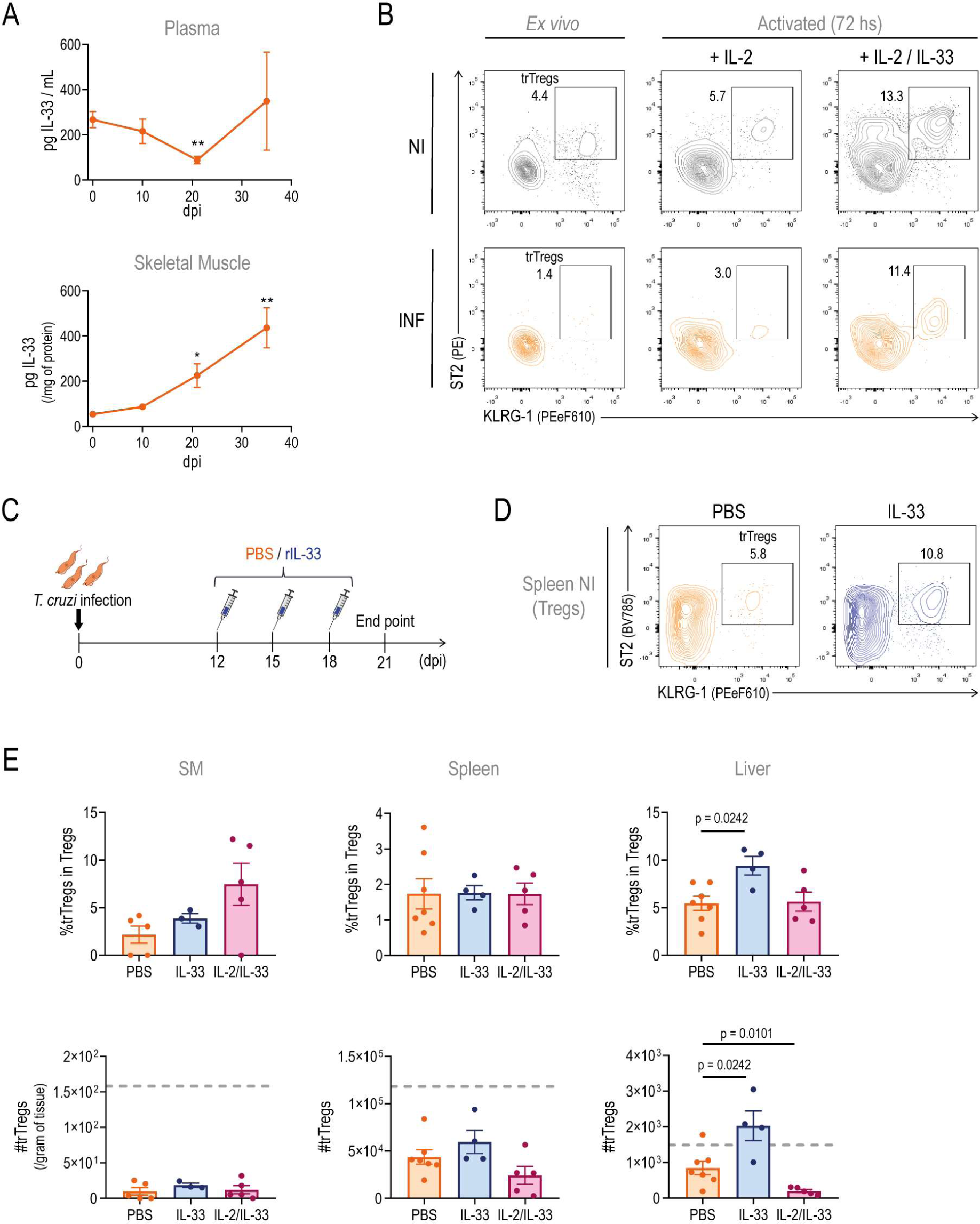
IL-33 supplementation fails to prevent trTregs reduction in established infection. (A) IL-33 concentration in plasma and SM lysates obtained from Foxp3-GFP mice at different days post infection (dpi). Muscle values were normalized to total protein content. Data are presented as mean ± SEM; N = 3-15 per dpi. (B) Representative dot plots showing ST2+ KLRG-1+ Tregs (trTregs) frequency within total Tregs isolated from the spleen of non-infected (NI) and infected (INF) (21 dpi) Foxp3-GFP mice. Left plots correspond to uncultured Tregs, while middle and right plots correspond to Tregs activated with anti-CD3+anti-CD28+IL-2 with or without rIL-33 for 72 hours. (C) Experimental scheme illustrating the treatment of Foxp3-GFP mice with an established infection with IL-33 or IL-33+IL-2. (D) Flow cytometry analysis of Tregs present in the spleen of non-infected (NI) Foxp3-GFP mice 72 h after receiving 3 doses of intraperitoneal IL-33 or PBS as described in (C). Dot plots show the frequency of trTregs within total Tregs. (E) Graphs displaying trTregs frequency within total Tregs (upper row) and absolute number (bottom row) in skeletal muscle (SM), liver and spleen from INF animals (21 dpi) receiving intraperitoneal PBS, IL-33 or IL-2/IL-33 as described in C. Bars indicate the mean ± SEM. For each tissue, gray dashed lines indicate the average of trTregs count in untreated NI mice. Statistical significance was determined by Kruskal-Wallis test (A) and Mann-Whitney test (E). P values in (A) are relative to 0 dpi: *p < 0.05; **p < 0.01; while in (E) represent pairwise comparison. Data are representative of two (A-B) and one (D-E) independent experiments.

### IL-33 supplementation expands trTregs from acutely infected animals *in vitro* but fails to prevent trTregs reduction in established infection

To investigate the possible role of trTregs in managing damage and the progression of pathology in *T. cruzi* infection, we aimed to increase trTregs numbers by administrating recombinant IL-33 (rIL-33) in our experimental setting. As a proof of principle, we first evaluated the responsiveness of trTregs from INF mice to IL-33. To this end, splenic Tregs (CD4+ Foxp3-GFP+) sorted from INF mice and NI mice were stimulated with anti-CD3, anti-CD28 and rIL-2 in the presence or absence of rIL-33 (Fig 3B). The addition of IL-33 to cultures containing Tregs from INF mice resulted in an expansion of trTregs, leading to a final percentage of ST2+ KLRG-1+ Tregs comparable to that obtained in IL-33-supplemented NI Tregs cultures. Notably, IL-33 supplementation failed to induce ST2 and KLRG-1 expression in conventional T cells (CD4+ Foxp3-GFP-) obtained from the spleens of either INF or NI animals (S4B Fig). These results indicate that rIL-33 specifically acts on the Tregs pool by expanding trTregs even when they are isolated from the *T. cruzi* infection environment, supporting its potential for treating INF animals.

For the *in vivo* treatment, with the aim of preventing the infection-induced reduction of trTregs, we administered rIL-33 (or PBS as control) intraperitoneally at 12, 15 and 18 dpi, as schematized in Fig 3C. In an alternative approach, mice were co-administered with rIL-2 and rIL-33 considering the relevance of IL-2 for Tregs survival [54] and taking into account that systemic levels of this cytokine remain unchanged even after the increased cell demand resulting from T cell expansion during acute *T. cruzi* infection [44]. As a positive control of its biological activity, we observed that *in vivo* intraperitoneal (i.p.) administration of IL-33 to NI animals led to trTregs expansion (Fig 3D), as previously reported [5]. Remarkably, the treatment with rIL-33 or rIL-33 plus rIL-2 in INF mice failed to increase trTregs frequency or absolute numbers in SM and spleen when compared to PBS-treated controls (Fig 3E). The injection of rIL-33 alone had effect on the liver, resulting in an increase in trTregs frequency and a subsequent recovery in trTregs absolute numbers, which reached levels similar to those observed in the livers of untreated NI mice (represented by grey dashed lines) (Fig 3E). To further address whether IL-33 supplementation had an impact on tissue damage or infection progression, despite the limited effect on trTregs, we initially examined the plasma levels of biochemical markers indicative of tissue damage. The treatment with rIL-33 did not improve these markers, while the injection of rIL-2 plus rIL-33 appeared to worsen them compared to PBS-treated controls (S4C Fig). Moreover, global indicators of disease progression, such as total weight loss at the peak of infection and overall survival, also showed no differences between treated and control mice (S4D and S4E Figs).

Since systemic (i.p.) treatment with IL-33 in INF mice did not yield any significant effects on the evaluated parameters, we speculated that local administration in peripheral tissue may better target trTregs. Therefore, we tested the effect of intramuscular (i.m.) rIL-33 injections. In this approach, INF mice were injected at 12, 15 and 18 dpi with rIL-33 (0.3μg per muscle) in one hind limb and with PBS in the other as control, according to the procedure described by Kuswanto et al., 2016 [12]. Similar to systemic treatment, i.m. rIL-33 injection in NI animals resulted in an accumulation of Tregs in SM, which contained a high proportion of trTregs (S4F Fig), while it had no effect on INF animals (S4G Fig).

Overall, these results indicate that, although trTregs from INF mice are intrinsically able to respond to IL-33, the reduction of trTregs that occurs during acute *T. cruzi* infection cannot be rescued by rIL-33 administration, likely due to particular signals generated in the context of an established infection.

### Inflammatory and microbial-derived signals are unable to restrict IL-33 mediated expansion of trTregs *in vitro*

To comprehend the mechanisms contributing to the limited impact of rIL-33 on inducing trTregs during *T. cruzi* infection, we directed our attention towards molecules known to counteract IL-33’s biological effects, potentially activated by this parasitic infection. Specifically, we examined soluble ST2 (sST2), a spliced variant of ST2 lacking the cytosolic and transmembrane domains, which acts as a decoy receptor to neutralize IL-33 activity under various inflammatory conditions [55]. Our findings revealed undetectable levels of sST2 in the serum from both NI and INF mice (Fig 4A). Within tissues, sST2 levels were shown to be similar between INF mice and NI counterparts in spleen and SM lysates while in the liver, sST2 levels could not be quantified as they were remarkably high, surpassing the methodological higher detection point even in diluted lysates.

**Fig 4:**
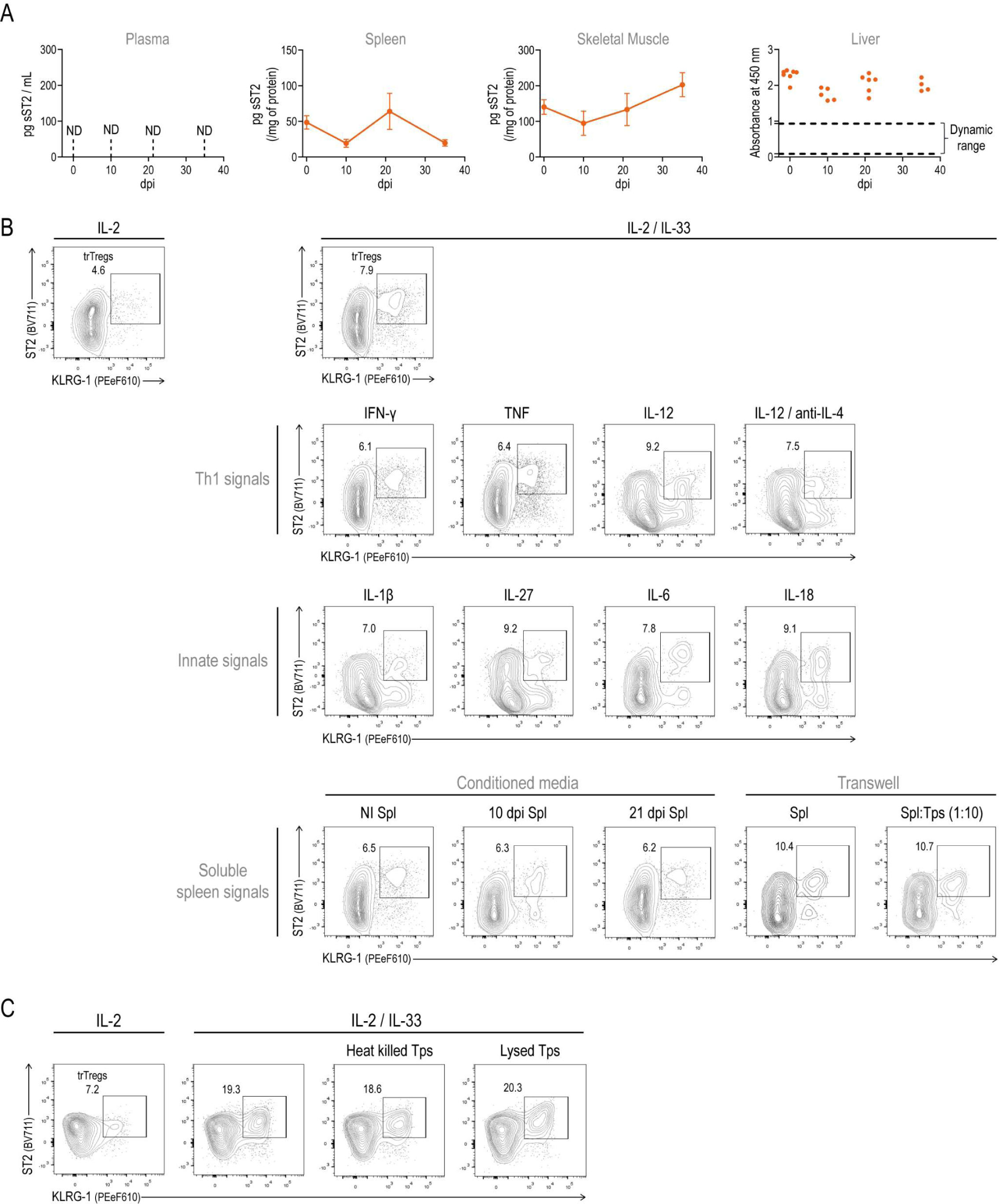
Inflammatory and microbial-derived signals are unable to restrict IL-33 mediated expansion of trTregs *in vitro*. (A) Soluble ST2 (sST2) concentration in plasma, as well as in spleen, skeletal muscle (SM) and liver lysates obtained from Foxp3-GFP mice at different days post infection (dpi). In spleen and SM, values were normalized to total protein content and are presented as mean ± SEM. In the liver, results from a 1/10 dilution of liver lysates are depicted as absorbance units due to falling outside the assay’s dynamic range, as indicated by the black dashed line. Statistical significance was determined by one-way ANOVA in spleen and SM. Representative of one experiment with N = 4-7 per dpi. ND: non-detectable. (B-C) Representative dot plots showing ST2+ KLRG-1+ Tregs (trTregs) frequency within total Tregs isolated from the spleen of non-infected (NI) Foxp3-GFP mice for 72 h with anti-CD3+anti-CD28 together with the addition of different cytokines as follow: IL-2; IL-2+IL-33 and IL2+IL33 plus: cytokines associated to Th1 signals, innate signals, as well as conditioned media or transwell co-cultures providing soluble spleen-derived signals (B) or microbial ligands (C), as indicated above each plot. (B-C) Data were collected from 4 independent experiments. Spl: splenocytes, Tps: trypomastigotes.

Beyond sST2, several pro-inflammatory cytokines such as IL-1β, IL-27, IFN-γ and TNF have been demonstrated to prevent IL-33 effects on target cells [56–58]. Indeed, the latter two can block *in vivo* the expansion of trTregs induced by rIL-33. Given the significant increase in some of these and other inflammatory signals like IL-6, IL-12 and IL-18 during acute *T. cruzi* infection [44,59,60], we explored their potential role in preventing the IL-33-mediated trTregs expansion in our experimental setting. To this end, we took advantage of the *in vitro* approach described in Fig 3B. Sorted splenic Tregs from NI mice were cultured in the presence of the trTregs expansion cocktail (anti-CD3, anti-CD28 plus rIL-2 and rIL-33) along with other recombinant cytokines, conditioned media or other stimuli (Fig 4B). None of the individual cytokines tested nor a Th1 differentiation cocktail (IL-12 plus anti-IL-4) were capable of preventing the expansion of trTregs induced by IL-33. We also considered that, in the context of *T. cruzi* infection, multiple inflammatory signals may concurrently exist and synergize to block IL-33’s effect. Therefore, we simulated these conditions using conditioned media obtained after 24-hour of polyclonal stimulation of splenocytes obtained from NI or INF (10 and 21 dpi) mice. None of these conditioned media were able to abrogate the IL-33-induced trTregs expansion. Then, a second approach involving transwell cultures was designed to emulate the early stages of infection. In this assay, where Tregs were co-cultured in the upper transwell chamber with splenocytes of NI mice either alone or in the presence of trypomastigotes, IL-2 and IL-33 also retained their capacity to expand trTregs.

Finally, considering that the maximal trTregs reduction correlates with the highest parasitemia, and taking into account previous results indicating that Treg differentiation is affected by the presence of parasites [44], we further evaluated a possible inhibitory effect from microbial ligands. To this end, heat-killed or lysed trypomastigotes were added to the cultures, however, even in the presence of these sources of microbial ligands, trTregs cell expanded normally in the presence of IL-2 + IL-33 (Fig 4C).

Collectively, these findings suggest that the ineffectiveness of IL-33 treatment in promoting trTregs cells is not likely due to increased levels of a well-known IL-33 modulator such as sST2 or the influence of prominent inflammatory cytokines or microbial ligands heightened in the context of *T. cruzi* infection. Instead, a complex combination of different signals, an unidentified modulator, or other mechanisms may underlie the absence of effect after IL-33 injection in established *T. cruzi* infection.

### Early rIL-33 administration expands trTregs and improve disease outcome in infected mice

In a subsequent effort to modulate the trTregs response *in vivo* during acute infection, and considering that results from the previous section suggested that IL-33 injection expands trTregs *in vivo* in NI mice but its effects are restrained by an unidentified signal(s) emerging along the infection, we opted to treat INF mice with rIL-33 on 0, 3, and 6 dpi (Fig 5A).

**Fig 5:**
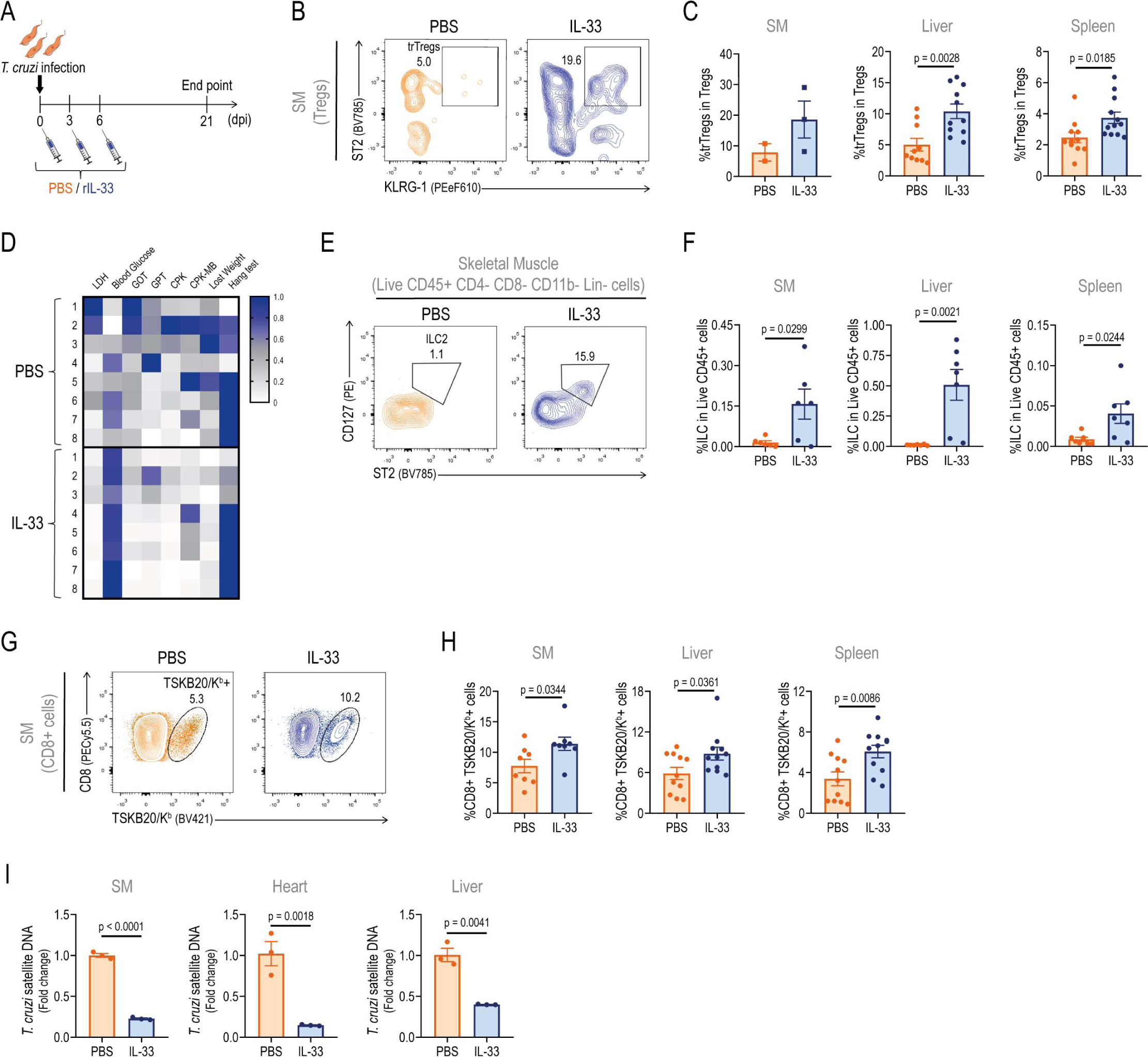
Early rIL-33 administration expands trTregs and improve disease outcome in infected mice. Immune response and disease progression was evaluated in infected Foxp3-GFP mice after receiving intraperitoneal IL-33 the day of infection and on 3 and 6 days post infection (dpi). (A) Experimental scheme. (B) Representative dot plots showing ST2+ KLRG-1+ Tregs (trTregs) frequency within total Tregs isolated from the skeletal muscle (SM) of PBS or IL-33-treated infected mice. (C) Graphs displaying trTregs frequency within total Tregs in SM, liver and spleen at 21 dpi. For SM, squares represent pools with N = 4-5. For spleen and liver, circles represent individual mice. (D) Heatmap showing relative values corresponding to parameters of disease progression at 21 dpi. Each column represents one mouse. (E) Representative dot plots illustrating type 2 innate lymphoid cells (ILC2) frequency within CD45+ CD4-CD8-Lin (CD3, CD19, NK1.1, CD11c)-CD11b-cells. Graphs correspond to SM from PBS or IL-33-treated infected mouse. (F) ILC2 frequencies within Live CD45+ cells in SM, liver and spleen at 21dpi. (G) Representative dot plots showing TSKB20/K^b^ staining in SM CD8+ cells from PBS or IL-33-treated infected mice. (H) Percentage of parasite-specific CD8+ T cells in SM, liver and spleen at 21dpi. (I) *T. cruzi* satellite DNA quantification in SM, heart and liver at 21 dpi. (C, F, H and I) Bars represent the mean ± SEM. Statistical significance was determined as follow: Mann-Whitney test for SM and unpaired t test for liver and spleen (A); unpaired t test (F and H) and Mann-Whitney test (I). Data are representative of two (C, D, H and I) and one (F) independent experiments.

Early IL-33 injection expanded trTregs in infected (INF) mice, as evidenced by the tendency for trTreg frequency and absolute numbers to increase in SM, alongside significant increases observed in the liver and spleen by 21 dpi (Figs 5B, 5C and S5A Fig). Of note, trTregs numbers from IL-33 treated INF animals exceeded those observed in untreated NI mice (S5A Fig, represented by grey dashed lines). These results illustrated the efficiency of early IL-33 treatment in sustaining elevated trTregs throughout at least the two-week period up to the peak of the infection, raising the possibility it may impact on tissue repair mechanisms and infection progression. To assess this, we measured several parameters such as biochemical markers of tissue damage, total body weight loss and SM strength (S5B-D Figs). The analysis of the outcome of these parameter evaluations revealed an overall improvement in the health state of the IL-33-treated group at 21 dpi, as summarized in the heatmap in Fig 5D. To characterize changes occurring in target tissues with IL-33 treatment, we performed histological analysis on SM. The results shown similar alterations in SM architecture between both experimental groups, characterized by the presence of a diffuse mononuclear infiltrate, necrosis and calcification of muscular fibers. Additionally, all mice showed scarce number of centrally nucleated muscle cells, which are associated with tissue regeneration [61] (S5E Fig).

The responsiveness to IL-33 is mediated by ST2, expressed in different immune cell types [16,55]. Therefore, it is expected that our early IL-33 treatment during *T. cruzi* infection may affect other ST2-expressing cell subsets involved in tissue damage control such as type 2 Innate lymphoid cells (ILC2) [49,62]. Accordingly, we assessed ILC2 infiltrate in different tissues using the gating strategy shown in S5F Fig, adapted from the protocol reported by Tait Wojno and Beamer [63]. As anticipated, ILC2 frequency and absolute numbers were also increased after rIL-33 injection during early *T. cruzi* infection (Figs 5E and 5F, and S5G Fig).

In addition to trTregs and ILC2, IL-33 can activate effector immune cells either directly through ST2 signaling on the target cell or indirectly by inducing the production of intermediate cues, such as IFN-γ [10,55,64,65]. We, therefore, evaluated if early IL-33 administration affected parasite specific CD8+ T cells, an effector response critical for *T. cruzi* control [66]. Interestingly, IL-33 treated animals showed increased frequencies of this cell subset in SM, liver as well as in spleen (Figs 5G and 5H). These changes in frequency corresponded with a tendency toward higher counts of parasite-specific CD8+ T cells in SM, conserved counts in the liver, and increased counts in the spleen (S5H Fig). In correlation with these results, INF mice that received IL-33 injection showed significantly decreased parasitism in non-lymphoid tissues such as SM, heart and liver (Fig 5I).

Altogether, these findings demonstrate that early rIL-33 administration can improve the course of acute infection, not only by reducing tissue damage, but also by increasing parasite control. These results may be attributed to the combined effect of IL-33 on different immune cell subsets such as trTregs, ILC2 and effector CD8+ T cells.

## DISCUSSION

Previous reports indicate that tissue injury, whether sterile or infection-derived, is associated with trTregs accumulation due to increased IL-33 release after cell destruction [16]. This reparative mechanism is well described in sterile damage, particularly within SM [4,11,12]. In the context of acute *T. cruzi* infection—a model for Chagas’ disease with a profound compromise of various target tissues, including SM—our results unveil a novel scenario marked by a diminished trTregs response and reduced plasmatic IL-33 concentration despite extensive tissue damage. This suggests a specific impairment in the regulatory fate associated with tissue repair, complementing our earlier work which reported limited Tregs responses but with the acquisition of a type 1 specialization profile during *T. cruzi* infection [44]. Altogether, our findings delineate a scenario where the reduction of a reparative/regulatory cell subset during the acute phase may potentially facilitate tissue damage by compromising physiological repair mechanisms. Indeed, a detailed characterization of tissue damage associated with *T. cruzi* infection, particularly in SM, revealed a clear modification in the SM transcriptome with activation of immune-related pathways and a general downregulation of genes associated with myogenesis, adipogenesis, and oxidative phosphorylation pathways, linked to normal muscle physiology and repair processes after injury [67–69]. Therefore, our results provide evidence that during acute *T. cruzi* infection, the transcriptional landscape in SM denotes damage and correlates with a reduced frequency of trTregs. This association may potentially connect deficient repair processes with long-term consequences in chronic immunopathology during Chagas’ disease. Further studies may be required to deeply assess these possible connections.

The attenuation of the reparative response during acute *T. cruzi* infection was systemic, manifested by reduced frequency and absolute numbers of trTregs across evaluated tissues, including SM, liver, and spleen. Unexpectedly, IL-33 levels exhibited both consistency and discrepancy: plasma IL-33 concentration diminished, correlating with the limited trTregs responses, while muscle IL-33 increased, yet failed to prevent the reduction of trTregs in that tissue. Two aspects of these findings were puzzling: the unexpected reduction in systemic IL-33 levels despite documented tissue damage in *T. cruzi* infection and the lack of an accompanying increase in trTregs in muscle despite an elevated local IL-33 concentration. The mechanisms regulating IL-33, which are complex and varied, include its rapid degradation after tissue injury [16,70,71]. For example, caspases 1 and 7, produced in inflammatory contexts and involved in the death of infected cells, fragment IL-33 to inactivate it [72]. Additionally, IL-33 is easily oxidizable in the extracellular space, limiting ST2-dependent immunological responses [73]. Accordingly, the inflammatory milieu during acute *T. cruzi* infection could rapidly oxidize and/or degrade IL-33, diminishing its availability, especially in plasma. In contrast, elevated IL-33 levels were detected in muscle. However, this quantification was conducted in tissue lysates, presenting technical limitations that impede confirmation of the extracellular presence and, consequently, the availability of IL-33 for ST2+ cells. Discrepancies between IL-33 concentration and trTregs numbers in SM suggest potential unavailability or counteraction by inflammatory signals. Alternatively, trTregs cells may face high mortality or reduced generation during the acute stage. Considering the multi-step development of trTregs beginning in the spleen [8,74], the reduction in IL-33 levels in this organ during acute infection could be linked to a lower generation of these cells and, consequently, their reduced arrival at target organs.

Given the scenario described earlier, and our aim to understand the impact of trTregs reduction on disease progression, we intended to boost the numbers of this cell subset through rIL-33 supplementation in established *T. cruzi* infection. However, IL-33 treatment, administered systemically or locally around the second week of infection, failed to rescue trTregs numbers and had no effect on disease progression. Notably, the lack of response to IL-33 in terms of trTregs expansion was specific to the infection condition, as this cytokine increased trTregs in non-infected animals. The ability of trTregs from infected mice to expand upon IL-33 stimulation *in vitro* suggests intrinsic responsiveness to this growth factor, pointing to a restrictive *in vivo* environment imposed by the infection. As we established that inflammatory environments or even the parasite itself might negatively affect Tregs differentiation [44], and considering that inflammatory cytokines can impede IL-33 signaling in trTregs [57], we investigated potential infection-associated cues that could counteract IL-33’s effect. We found that sST2, a natural decoy receptor of IL-33 signaling [55], is not increased during *T. cruzi* infection, suggesting that it is unlikely involved in the lack of response to IL-33 in our infection setting. Further evaluation through *in vitro* approaches, combining rIL-33 with various inflammatory or parasite signals, failed to identify any molecule, cocktail, or conditioned media capable of blocking IL-33-mediated expansion of trTregs. These results emphasize that the intricate combination of signals present in a living host undergoing acute *T. cruzi* infection is challenging to replicate with *in vitro* approaches, making it difficult to identify critical determinants of an environment particularly restrictive for regulatory pathways.

As an alternative strategy to assess IL-33 effects while avoiding the restrictive environment around the peak of *T. cruzi* infection, we implemented early treatment during the first week of infection. This approach resulted in a less severe acute infection progression along with better parasite control. Given that IL-33 is able to modulate several immune cell subsets [70], we focused not only on trTregs but also on ILC2 and CD8+ T cells. Early IL-33 supplementation during *T. cruzi* infection led to a significant expansion of trTregs, consistent with previous reports [12,20,24]. Notably, the magnitude of the trTregs response increased not only in secondary lymphoid organs such as the spleen but also in target organs like the liver and SM, remaining evident even at the infection’s peak. Interestingly, early IL-33 treatment also elevated ILC2 and parasite-specific CD8+ T cell frequencies. ILC2 play crucial roles in combating certain infectious agents and, similar to trTregs, promote tissue reparative processes [62]. Indeed, IL-33-mediated ILC2 expansion has been linked to infection resistance against cerebral malaria and various intestinal infections [23,75–77]. Moreover, our comprehensive evaluation of immune cell subsets highlighted that IL-33 potentiated antiparasitic CD8+ T cell immunity, aligning with previous research that suggested a role for this alarmin in inducing robust antiviral responses [78,79]. In particular, IL-33 has been recently shown to preserve CD8+ T cell stemness and re-expansion capacity in the context of a chronic viral infection [80].

In summary, our findings underscore the positive impact of IL-33 supplementation during early *T. cruzi* infection, contributing to reduced tissue damage and improved parasite control. Building upon existing research, we propose that this beneficial effect arises from the concerted action of expanding cell subsets involved in tissue repair, such as trTregs and ILC2, alongside those engaged in microbial control, particularly CD8+ T cells. Future studies are warranted to meticulously dissect the specific roles of each of these immune cell subsets in mediating the effects of IL-33 on *T. cruzi* infection outcome. Interestingly, the simultaneous expansion of trTregs and parasite-specific CD8+ T cells might seem counterintuitive, given numerous reports linking regulatory responses with diminished antimicrobial immunity [81]. Our previous study in the context of *T. cruzi* infection demonstrated that adoptive Tregs transfer represses parasite-specific CD8+ T cell responses [44]. Furthermore, it has been shown that tissue resident Tregs exhibit heightened regulatory features [25]. Significantly, our results challenge this notion by revealing that boosting specialized regulatory T cell responses does not necessarily attenuate effector responses. This simultaneous enhancement of both regulatory/reparative and antiparasitic cell subsets, likely a result of the individual effects of IL-33 on each ST2-expressing subset, proves to be beneficial for the progression of acute *T. cruzi* infection. This discovery establishes a precedent, suggesting the potential for rational novel therapies in Chagas’ Disease or other infectious diseases. To date, only one study has evaluated IL-33 expression in chronic Chagas disease patients showing no correlation with disease severity [82]. Therapies involving IL-33 could be designed to modulate the immune system, favoring a specific combination of regulatory and effector responses. The goal would be to enable effective pathogen clearance while minimizing collateral damage, thereby preventing clinical pathology.

## MATERIALS AND METHODS

### Ethics statement

Mouse handling followed international ethical guidelines. All experimental procedures were conducted in compliance with the ethical standards set by the Institutional Animal Care and Use Committee of Facultad de Ciencias Químicas – Universidad Nacional de Córdoba, and were approved under protocol numbers RD-733-2018.

### Mice

Age-matched (8 to 12 week-old) mice of both sexes were used. Foxp3-GFP reporter mice (B6.Cg-Foxp3tm2Tch/J) were obtained from The Jackson Laboratories (USA). BALB/c mice were obtained from School of Veterinary, La Plata National University (La Plata, Argentina). Animals were bred in the animal facility of the Facultad de Ciencias Químicas, Universidad Nacional de Córdoba, and housed under a 12:12 h light-dark cycle with food and water ad libitum. The institutional animal facility follows the recommendations of the Guide for the Care and Use of Experimental Animals, published by the Canadian Council for the Protection of Animals.

### Parasites and experimental infection

For all experiments, *T. cruzi* Tulahuén strain was used. Bloodstream trypomastigotes were maintained in male BALB/c mice by serial passages every 10-11 days. For *in vivo* assays, Foxp3-GFP reporter mice were inoculated intraperitoneally with 0.2 mL PBS containing 5000 trypomastigotes.

For *in vitro* assays, trypomastigotes were obtained from the extracellular medium of infected monolayers of Vero cells cultured in RPMI 1640 medium (Gibco, Invitrogen) containing 10% heat inactivated fetal bovine serum (FBS, Natocor), 2 mM glutamine (Gibco), 10 mM HEPES (Gibco) and 40 ug/mL gentamicin (Veinfar Laboratories). After 7 days of infection, extracellular medium was collected, centrifuged at 1800 g for 30 min at room temperature and incubated for 2 h at 37 °C. Trypomastigotes were recovered from the supernatant and counted using a Neubauer chamber. Heat-killed trypomastigotes were obtained after incubation at 56 °C for 10 min (adapted from [83]), while lysed trypomastigotes were obtained after 3 cycles of freeze/thaw and 5 minutes of sonication (adapted from [84]).

### Parasite quantification in blood and tissues

Parasitemia was monitored by counting the number of viable trypomastigotes in blood after lysis with a 0.87% ammonium chloride buffer. For tissue parasite quantification, genomic DNA was purified from 50 μg of tissue (spleen, liver, SM and heart) using TRIzol Reagent (Life Technologies) following manufactureŕs instructions. Satellite DNA from *T. cruzi* (GenBank AY520036) was quantified by real time PCR using specific Custom Taqman Gene Expression Assay (Applied Biosystem) using the primer and probe sequences described by Piron et al. [85]. The samples were subjected to 45 PCR cycles in a thermocycler StepOnePlus Real-Time PCR System (Applied Biosystems). Abundance of satellite DNA from *T. cruzi* was normalized to the abundance of GAPDH (Taqman Rodent GAPDH Control Reagent, Applied Biosystem), quantified through the comparative CT method and expressed as arbitrary units, as previously reported [44,47,86].

### Cell preparation

To obtain cell suspensions from solid tissues, euthanized mice were perfused with 10 mL cold Hanks’ Balanced Salt Solution (Gibco). Spleens and livers were obtained and mashed through a tissue strainer. Liver infiltrating cells were obtained after 25 min centrifugation (600 g without brake) in a 35% and 67.5% bilayer Percoll (GE Healthcare) gradient. The interphase containing leukocytes was recovered and washed. Erythrocytes in spleen and liver cell suspensions were lysed for 3 min in ACK Lysing Buffer (Gibco). Heart and SM (quadriceps, gastrocnemius and tibialis anterior) were excised, minced and digested for 30 min in collagenase D (2 mg/mL, Roche) and DNase I (100 μg/mL, Sigma). Digested tissues were filtered through a 70 μm filter and washed. Infiltrating leucocytes were obtained after 25 min centrifugation (600 g without brake) in a 40% and 75% bilayer Percoll gradient. The interphase was recovered and washed. Cell numbers were counted in Turk’s solution using a Neubauer chamber.

### In vitro assays

For Tregs and Tconv culture, cells were purified from NI or 21 dpi Foxp3-GFP mice. CD4+ cells were isolated from pooled splenic suspensions by magnetic negative selection using EasySep Mouse CD4+ T Cell Isolation Kit (StemCell Technologies) according to manufacturer’s instruction. Afterwards, the enriched CD4+ T cell suspension was surface stained and Tregs and Tconv were further purified by cell sorting with a FACSAria II (BD Biosciences) according to the following phenotype: Tregs (CD4+ Foxp3-GFP+) and Tconv (CD4+ Foxp3-GFP-). Purified cells (75000 cells/well) were cultured for 3 days in 96-well U bottom plates coated with 2 μg/mL anti-CD3 (eBioscience) and 1 μg/mL anti-CD28 (eBioscience) supplemented with 10ng/mL of recombinant mIL-2 (Biolegend) to allow Treg survival. Cells were cultured in complete culture media containing RPMI 1640 medium (Gibco, Invitrogen) 10% heat inactivated FBS (Natocor), 2mM glutamine (Gibco, Invitrogen), 55uM 2-mercaptoethanol (Gibco, Invitrogen) and 80ug/mL gentamicin (Veinfar Laboratories). When indicated, media contained 50 ng/mL recombinant mIL-33 (R&D) alone or combined with the following murine recombinant cytokines: IFN-γ (50 ng/mL, Immunotools), TNF (50 ng/mL, Immunotools), IL-1β (2 ng/mL, R&D), IL-6 (20 ng/mL, Shenandoah), IL-12p70 (10 ng/mL, Peprotech), IL-18 (50 ng/mL, R&D), IL-27 (20 ng/mL, R&D) as well as with 2 μg/mL anti-mouse IL-4 (Peprotech). Alternatively, 50 μL of conditioned media was used.

In co-culture transwell experiments, Tregs from NI mice were placed in the bottom of the culture plate in the presence of recombinant mIL-33 and splenocytes (1:1 ratio) alone or with trypomastigotes (1:10 ratio) that were placed in the transwell chamber.

In co-cultures with parasite ligands, Tregs from NI animals were incubated with heat-killed or lysed trypomastigotes (ratio 1:1) in the presence of recombinant mIL-33.

### Conditioned media generation

For conditioned media, total splenocytes were isolated from pooled splenic suspensions of NI, 10 dpi and 21 dpi Foxp3-GFP mice. Cell suspensions (5×10^6^ cells/ml) were cultured for 24 h in 24-well plates in complete culture media supplemented with 50 ng/mL PMA and 1 μg/mL ionomycin (Sigma-Aldrich).

### Biochemical determinations

Plasma was collected after blood centrifugation for 8 min at 3000rpm. Quantification of biochemical markers of tissue damage was performed at Laboratorio Biocon (Córdoba, Argentina) using a Dimension RXL Siemens analyzer. GOT, GPT, LDH and CPK activity was determined by UV kinetic method, CPK-MB activity was evaluated by enzymatic method, while glucose concentration was assessed by kinetic/colorimetric method.

### IL-33 and sST2 quantification

IL-33 concentration was determined with an IL-33 Mouse ELISA kit (eBioscience), while sST2 was quantified using a Mouse ST2/IL-33R DuoSet ELISA kit (R&D Systems) in plasma and tissue lysates. Plasma samples were obtained as previously described. Tissue lysates were obtained after centrifugation at 10000g during 10 min of tissue samples homogenized in PBS containing 0,5% BSA, 0,4 M NaCl, 1 mM EDTA, 0,05% Tween 20 and a protease inhibitor cocktail (Sigma-Aldrich) (adapted from [87]). GraphPad Prism 8.0.1 software was used to generate the calibration curve and determine IL-33 and sST2 concentration. In tissue lysates, values were normalized to total protein content determined using Bradford reagent (BioRad). Two Synergy HT Multi-mode microplate reader (Biotek) was used to determine absorbances at 450 nm (ELISA) and 595 nm (protein quantification).

### Flow cytometry

Combinations of the following antibodies were used for flow cytometry: biotin polyclonal anti-Amphiregulin (R&D Systems), PE anti-BATF clone S39-1060 (BD Pharmingen), Super Bright 645 anti-CD11b clone M1/70 (eBioscience), PE-Cyanine7 anti-CD11c clone N418 (eBioscience), PE anti CD127 clone A7R34 (eBioscience), PE-Cyanine7 anti-CD19 clone eBio1D3 (eBioscience), PE-Cyanine7 anti-CD3 clone 145-2C11 (eBioscience), APC, Super Bright 645 and APC-eFluor 780 anti-CD4 clone GK1.5 (eBioscience), PE-Cyanine5 anti-CD44 clone IM7 (eBioscience), Alexa Fluor 700 and APC-Cyanine7 anti-CD45 clone 30-F11 (eBioscience and BD Pharmingen respectively), APC-eFluor 780 anti-CD62L clone MEL-14 (eBioscience), PE-Cyanine5.5 anti-CD8 clone 53-6.7 (eBioscience), Brilliant Violet 605 anti CTLA-4 clone UC10-4B9 (Biolegend), FITC anti-Foxp3 clone FJK-16s (eBioscience), Super Bright 600 anti-GITR clone DTA-1 (eBioscience), PerCp-eFluor 710 anti-ICOS clone 7E.17G9 (eBioscience), PerCp-eFluor 710 anti-IRF4 clone 3E4 (eBioscience), eFluor 660 anti-Ki-67 clone SolA15 (eBioscience), PE-eFluor 610 anti-KLRG-1 clone 2F1 (eBioscience), PE-Cyanine7 anti-NK1.1 clone PK136 (Biolegend), Brilliant Violet 421 anti-PD-1 clone 29F.1A12 (Biolegend), PE and Brilliant Violet 421 anti-ST2 clone DIH9 (Biolegend) and PerCp-eFluor 710 anti-TIGIT (eBioscience). To detect biotinylated antibodies, Streptavidin Qdot 605 (Invitrogen) was used.

For surface staining, cell suspensions were incubated with fluorochrome labeled-antibodies together with LIVE/DEAD Fixable Aqua Dead Cell Stain Kit, for 405 nm excitation (Invitrogen) in PBS 2% FBS for 20 min at 4°C. To identify *T. cruzi* specific CD8+ T cells, cell suspensions were incubated with an H-2Kb *T. cruzi* trans-sialidase amino acids 569-576 ANYKFTLV (TSKB20) APC-or Brilliant Violet 421-labeled Tetramer (NIH Tetramer Core Facility) for 20 min at 4 °C, in addition to the surface staining antibodies.

For transcription factors detection, cells were initially stained for surface markers, washed, fixed, permeabilized and stained with Foxp3/Transcription Factor Staining Buffers (eBioscience) according to eBioscience One-step protocol for intracellular (nuclear) proteins. For intracellular cytokine detection, 2 x 10^6^ cells per well were cultured in 200 μL supplemented RPMI 1640 medium and stimulated during 2 h at 37 °C with 50 ng/mL PMA and 1 µg/mL ionomycin (Sigma-Aldrich) in the presence of Brefeldin A and Monensin (eBioscience). Then, stimulated cells were surface-stained as indicated above, fixed and permeabilized with Intracellular Fixation & Permeabilization Buffer Set (eBioscience) or IC Fixation Buffer and permeabilization Buffer (eBioscience) following manufacturers’ indications. In all cases, intracellular staining was performed by a 30 min incubation at room temperature.

All samples were acquired on FACSCanto II (BD Biosciences), LSRFortessa (BD Biosciences) or Attune-NxT (Life Technologies) and data were analyzed with FlowJo software version X.0.7. For cell sorting, FACS Aria II (BD Biosciences) was used.

### RNA sequencing

Perfused mouse quadriceps were obtained and stored in RNAlater Stabilization Solution (Invitrogen) at −80°C. Then, 25mg of tissue was dissociated using Bead Ruptor Elite (Omni International) for 45 seconds at 4.85m/s and RNA was isolated using RNeasy Fibrous Tissue Mini Kit 50 (Qiagen) following manufacturer’s indications. RNA concentration was determined with a Qubit 2.0 Fluorometer (Invitrogen), while a 2100 Bioanalyzer (Agilent) was used for quality evaluation. Poly(A) mRNA Magnetic Isolation Module (New England BioLabs) was used for cDNA library preparation according to manufacturer’s protocol. Quality control of libraries was determined as described for RNA. Quantification of cDNA libraries was performed with PerfeCTa NGS Quantification Kit for Illumina Sequencing Platforms (QuantaBio). For each experimental group, 3 biological replicates were sequenced with NextsSeq 550 (Illumina). For data analysis, a Salmon index was built from Gencode [88] Mouse release M27 (GRCm39) [GENCODE - Mouse Release M27 (gencodegenes.org)] using Salmon [89] v1.5.1 [Release Salmon 1.5.1 · COMBINE-lab/salmon · GitHub] with default k-mer size (31) and the--gencode flag. FASTQ sequence reads (SRA accession PRJNA941341) were mapped to the M27 index and transcript abundances were estimated using salmon quant on 8 threads. Salmon quant files were subsequently loaded into R v4.1.0 using tximeta v1.10.0, and differentially expressed genes were called using default parameters in DESeq2 v1.32.0 per the Bioconductor vignette [Analyzing RNA-seq data with DESeq2 (bioconductor.org)]. Genes with adjusted p-values ≤ 0.05 and |log_2_ fold-change| ≥ 1 were considered differentially-expressed and included in downstream analysis. For gene set enrichment analysis, EnrichR tool [90–92] was used an d MSigDB Hallmark 2020 gene sets were interrogated. Volcano plots were generated using “ggplot2” package in R. The datasets generated for this study can be found in the NIH repository under accession number PRJNA941341 (https://www.ncbi.nlm.nih.gov/sra/PRJNA941341).

### *In vivo* rIL-33 and rIL-2 treatment

Recombinant mIL-33 (Shenandoah) was administered via i.p. (2 μg in a total volume of 200 μL) or i.m. (0.3 μg/muscle in a total volume of 30-50 μL) at the specified time points. Dose and frequency of injections were adapted from reported protocols [12,20,24]. For i.m. treatment, quadriceps, gastrocnemius and tibialis anterior from the same hindlimb received each the dose detailed above. When indicated, i.p. injections also contained 1 μg of recombinant murine IL-2 (Gibco). PBS was used as vehicle.

### Muscle strength and body weight evaluation

Muscle strength was assessed by Kondziela’s inverted screen test (hang test) [93]. Mice were kept in the experimental room for 20 minutes before the test to ensure proper adaptation to the environment. Each mouse was placed in the center of a 43 cm^2^ wire mesh consisting of 12 mm squares of 1 mm diameter wire and surrounded by a 4 cm deep wooden frame. Screen was inverted and time was measured until the mouse fell off. Maximum test duration was 2 minutes. Total body weight was determined using a precision laboratory balance.

### Histological analysis

Perfused mouse quadriceps were fixed in formaldehyde solution and embedded in paraffin. Five μm thick sections were stained with activated hematoxylin followed by eosin alcoholic solution. Histopathological evaluation was performed by a pathologist under light microscopy. Photographs were taken using a Nikon Eclipse TE 2000 U equipped with a digital video camera.

### Statistics and graph creation

Unless otherwise indicated, both statistics calculation and graphs creation were performed with GraphPad Prism 8.0.1 software. The normality of data distribution was assessed using Shapiro-Wilk normality test. Statistical significance of mean value comparisons was determined using t-test or One-way ANOVA for normally distributed data, and Mann-Whitney test or Kruskal-Wallis test for non-normally distributed data, as appropriate. P values ≤ 0.05 were considered statistically significant and are indicated in the graphs. Outliers were identified using the ROUT method. Data are presented as mean ± SEM and the number of animals of each experimental group is indicated in the figure legends or shown in the plots. Principal Component Analysis graph and volcano plots were generated using “ggplot2” package in R software. Flow cytometry plots were exported from FlowJo software version X.0.7 after data analysis.

### AI Language Model Assistance

We used ChatGPT (developed by OpenAI) to assist in refining the written content of this study. ChatGPT provided suggestions and corrections based on the input provided by the user, enhancing the clarity and grammar of the text. ChatGPT output was critically revised by the user to ensure it conveys the desired message

## ACKNOWLEDGMENTS

We thank M. P. Abadie, M. P. Crespo, V. Blanco, D. Lutti, C. Noriega, F. A. Frontera, S. R. Oms, R. E. Villarreal, G. Furlán, N. M. Maldonado, A. Romero, L. V. Gatica and M. S. Miró (Centro de Investigaciones en Bioquímica Clínica e Inmunología) for their excellent technical assistance. We acknowledge the NIH Tetramer Core Facility for provision of APC- and Brilliant Violet 421-labeled TSKB20 tetramers. We thank S. Sandrone (Hospital Rawson, Córdoba, Argentina) for her professional commitment regarding histopathological analysis. We are also grateful to S. B. Lakshminarayana, C. Osborn, D. Kristen, D. Patra for technical assistance and J. Spector (BioMedical Research, Novartis, United States) for coordinating the global health fellowship.

## AUTHOR CONTRIBUTIONS

**Conceptualization:** Eva V. Acosta Rodríguez.

**Formal Analysis:** Santiago Boccardo.

**Funding Acquisition:** Eva V. Acosta Rodríguez.

**Investigation:** <u>All experimental procedures but RNAseq</u>: Santiago Boccardo (sample collection and processing and data analysis), Constanza Rodriguez, Cintia L. Araujo Furlan, Carolina P. Abrate, Laura Almada, Camila M. S. Giménez (sample collection and processing).

<u>RNAseq study:</u> Santiago Boccardo and Manuel A. Saldivia Concepción (sample collection and processing), Peter Skewes-Cox (data compilation), Srinivasa P. S. Rao (supervision and funding).

**Methodology:** Santiago Boccardo, Eva V. Acosta Rodríguez.

**Project Administration:** Eva V. Acosta Rodríguez, Adriana Gruppi, Carolina L. Montes. **Resources:** Eva V. Acosta Rodríguez, Adriana Gruppi, Carolina L. Montes, Srinivasa P. S. Rao. **Supervision:** Eva V. Acosta Rodríguez.

**Validation:** Santiago Boccardo, Eva V Acosta Rodríguez.

**Visualization:** Santiago Boccardo.

**Writing – Original Draft Preparation:** Santiago Boccardo, Eva V Acosta Rodríguez.

**Writing – Review & Editing:** Adriana Gruppi, Carolina L Montes, Constanza Rodriguez, Cintia L. Araujo Furlan, Carolina P. Abrate, Laura Almada, Camila M. S. Giménez, Manuel A. Saldivia Concepción, Peter Skewes-Cox, Srinivasa P. S. Rao.

## SUPPORTING INFORMATION

**S1 Fig: Peripheral target tissues display inflammatory responses during acute *T. cruzi* infection.** Inflammatory response was evaluated in *T. cruzi* infected Foxp3-GFP mice at different days post infection (dpi). (A) Whole quadriceps muscle (SM) RNAseq data analysis from NI and INF mice as described in Fig 1F-G; N = 3 per group. Volcano plots displaying differentially expressed genes (dots) between INF SM and NI SM. According to Figure 1F, genes associated with interferon gamma response, interferon alpha response and complement pathways are highlighted in red. (B) Number of CD45+ cells in heart and liver, determined by flow cytometry at different dpi. Data are presented as mean ± SEM; N = 4-9 per group. For liver, counts correspond to total leukocyte numbers, whereas for heart, cell numbers are normalized to tissue weight. Statistical significance was determined by one-way ANOVA and P values relative to 0 dpi. *p < 0.05; ****p < 0.0001.

**S2 Fig: Tregs frequency is reduced in lymphoid and non-lymphoid target tissues during acute *T. cruzi* infection.** Tregs response was evaluated by flow cytometry in spleen, liver, skeletal muscle (SM) and heart from non-infected (NI) and infected (INF) (21 days post infection) Foxp3-GFP mice. (A) Representative dot plots showing the frequency of Tregs (CD4+ Foxp3-GFP+) within total CD4+ cells from each tissue. (B) Bars displaying Tregs frequency within total CD4+ cells as the mean ± SEM. Circles represent individual mice; squares represent pools with 3-5 mice. For SM and heart, cell counts are normalized to tissue weight. Statistical significance was determined by unpaired t test for spleen, liver and SM; and by Mann-Whitney test for heart. P values are indicated in the graphs. (A-B) Data were collected from 3 independent experiments.

**S3 Fig: ST2+ KLRG-1+ Tregs from *T. cruzi* infected mice exhibit a phenotype compatible with *bona fide* trTregs.** Flow cytometry phenotypic analysis of Tregs subsets present in the spleen of non-infected (NI) or infected (INF) (21 dpi) Foxp3-GFP mice. Histograms show the expression of each cell marker in ST2+ KLRG-1+ (pink) and ST2-KLRG-1-(golden) Tregs as defined in Fig 2A. Numbers on top right corner of each plot indicate either mean fluorescence intensity or frequency of positive cells for each marker.

**S4 Fig: IL-33 supplementation fails to prevent trTregs reduction in established *T. cruzi* infection.** (A) IL-33 concentration was evaluated in spleen and liver lysates obtained from Foxp3-GFP mice at different days post infection (dpi). Values were normalized to total protein content. Data are presented as mean ± SEM; N = 4-12 per dpi. (B) GFPneg CD4+ conventional T cells (Tconv) isolated from the spleen of non-infected (NI) and infected (INF) (21 dpi) Foxp3-GFP mice were evaluated by flow cytometry. Representative dot plots showing ST2+ KLRG-1+ cells frequency within total Tconv. Left plots correspond to uncultured Tconv, while middle and right plots correspond to Tconv activated with anti-CD3+anti-CD23+IL-2 with or without rIL-33 for 72 hours. (C-E) Analysis of disease progression in INF mice treated with intraperitoneal IL-33 or IL-33+IL-2 as described in Fig 3C. (C) Plasma LDH, GOT, GPT, CPK and CPK-MB activities, and glucose concentration at 21 dpi. (D) Percentage of total body weight reduction at 21 dpi compared to 15 dpi. (E) Survival curve in the different experimental groups. (F, G) Representative plots depicting the frequencies of GFP+ CD4+ Tregs cells (F, G) and ST2+ KLRG-1+ cells (G) in NI (F) and INF (G) mice receiving intramuscular IL-33. Statistical significance was determined by Kruskal-Wallis test (A), Mann-Whitney test (C-D), Mantel-Cox test (E) and Wilcoxon test (G). P values in (A) are relative to 0 dpi: ****p < 0.0001; while in (C-D and G) represent pairwise comparisons; ns: non-significant.

**S5 Fig: Early rIL-33 administration improves the global health status without reducing SM alterations during acute infection.** Immune response and disease progression was evaluated in infected Foxp3-GFP mice after receiving intraperitoneal IL-33 the day of infection and on 3 and 6 days post infection (dpi). (A) trTregs count in skeletal muscle (SM), liver and spleen at 21 dpi. For SM, squares represent pools with N=4-5. For spleen and liver, circles represent individual mice. For each tissue, gray dashed lines indicate the average of trTregs count in untreated non-infected (NI) mice. (B) Plasma LDH, GOT, CPK, GPT and CPK-MB activities, and glucose concentration at 21 dpi. (C) Total body weight loss between 15 and 21 dpi. (D) Inverted screen test (max: 120 seconds) and (E) Hematoxylin-Eosin stain of quadriceps muscle at 21 dpi. Images are representative of N = 4. Black arrow: centrally nucleated muscular fibers. Magnification = 4X (left) and 10X (right). (F) Gating strategy used to identify type 2 innate lymphoid cells (ILC2) as CD45+ CD4-CD8-Lin (CD3, CD19, NK1.1, CD11c)-CD11b-CD127+ ST2+ cells. Dot plots are representative of SM from IL-33-treated infected mouse. (G) ILC2 and (H) parasite-specific CD8+ cell count in SM, liver and spleen at 21 dpi. For SM, counts are normalized to tissue weight. (A-D, G and H) Bars represent the mean ± SEM. Statistical significance was determined as follow: Mann-Whitney test for SM and unpaired t test for liver and spleen (A); unpaired t test (B-D, G and H). (A-D, G and H) P values from pairwise comparison are indicated in the graphs. Data are representative of two (A-D and H) and one (E and G) independent experiments.

**S1 Table: List of genes associated with inflammatory response and skeletal muscle physiology.** Differentially expressed genes between infected (15 days post infection) and non-infected mice are shown for each cellular pathway.

## Notes

### Competing Interest Statement

The other authors have declared no competing interest.
Manuel A. Saldivia Concepcion, Peter Skewes-Cox and Srinivasa P. S. Rao are employees from BioMedical Research, Novartis.

## REFERENCES

1. Zhao H, Liao X, Kang Y. Tregs: Where We Are and What Comes Next? Front Immunol. 2017;8: 1578. doi:10.3389/fimmu.2017.01578

2. Sakaguchi S, Mikami N, Wing JB, Tanaka A, Ichiyama K, Ohkura N. Regulatory T Cells and Human Disease. Annu Rev Immunol. 2020;38: 541–566. doi:10.1146/annurev-immunol-042718-041717

3. Cretney E, Kallies A, Nutt SL. Differentiation and function of Foxp3(+) effector regulatory T cells. Trends Immunol. 2013;34: 74–80. doi:10.1016/j.it.2012.11.002

4. Panduro M, Benoist C, Mathis D. Tissue Tregs. Annu Rev Immunol. 2016;34: 609–633. doi:10.1146/annurev-immunol-032712-095948

5. Delacher M, Imbusch CD, Weichenhan D, Breiling A, Hotz-Wagenblatt A, Träger U, et al. Genome-wide DNA-methylation landscape defines specialization of regulatory T cells in tissues. Nat Immunol. 2017;18: 1160–1172. doi:10.1038/ni.3799

6. Cho J, Kuswanto W, Benoist C, Mathis D. T cell receptor specificity drives accumulation of a reparative population of regulatory T cells within acutely injured skeletal muscle. Proc Natl Acad Sci U S A. 2019; 201914848. doi:10.1073/pnas.1914848116

7. Delacher M, Imbusch CD, Hotz-Wagenblatt A, Mallm J-P, Bauer K, Simon M, et al. Precursors for Nonlymphoid-Tissue Treg Cells Reside in Secondary Lymphoid Organs and Are Programmed by the Transcription Factor BATF. Immunity. 2020;52: 295–312.e11. doi:10.1016/j.immuni.2019.12.002

8. Li C, DiSpirito JR, Zemmour D, Spallanzani RG, Kuswanto W, Benoist C, et al. TCR Transgenic Mice Reveal Stepwise, Multi-site Acquisition of the Distinctive Fat-Treg Phenotype. Cell. 2018;174: 285–299.e12. doi:10.1016/j.cell.2018.05.004

9. Cipolletta D, Feuerer M, Li A, Kamei N, Lee J, Shoelson SE, et al. PPAR-γ is a major driver of the accumulation and phenotype of adipose tissue Treg cells. Nature. 2012;486: 549–553. doi:10.1038/nature11132

10. Schiering C, Krausgruber T, Chomka A, Fröhlich A, Adelmann K, Wohlfert EA, et al. The alarmin IL-33 promotes regulatory T-cell function in the intestine. Nature. 2014;513: 564–568. doi:10.1038/nature13577

11. Burzyn D, Kuswanto W, Kolodin D, Shadrach JL, Cerletti M, Jang Y, et al. A special population of regulatory T cells potentiates muscle repair. Cell. 2013;155: 1282–1295. doi:10.1016/j.cell.2013.10.054

12. Kuswanto W, Burzyn D, Panduro M, Wang KK, Jang YC, Wagers AJ, et al. Poor Repair of Skeletal Muscle in Aging Mice Reflects a Defect in Local, Interleukin-33-Dependent Accumulation of Regulatory T Cells. Immunity. 2016;44: 355–367. doi:10.1016/j.immuni.2016.01.009

13. Ali N, Zirak B, Rodriguez RS, Pauli ML, Truong H-A, Lai K, et al. Regulatory T Cells in Skin Facilitate Epithelial Stem Cell Differentiation. Cell. 2017;169: 1119–1129.e11. doi:10.1016/j.cell.2017.05.002

14. Nosbaum A, Prevel N, Truong H-A, Mehta P, Ettinger M, Scharschmidt TC, et al. Cutting Edge: Regulatory T Cells Facilitate Cutaneous Wound Healing. J Immunol. 2016;196: 2010–2014. doi:10.4049/jimmunol.1502139

15. Dombrowski Y, O’Hagan T, Dittmer M, Penalva R, Mayoral SR, Bankhead P, et al. Regulatory T cells promote myelin regeneration in the central nervous system. Nat Neurosci. 2017;20: 674–680. doi:10.1038/nn.4528

16. Cayrol C, Girard J-P. Interleukin-33 (IL-33): A nuclear cytokine from the IL-1 family. Immunol Rev. 2018;281: 154–168. doi:10.1111/imr.12619

17. Astarita JL, Dominguez CX, Tan C, Guillen J, Pauli ML, Labastida R, et al. Treg specialization and functions beyond immune suppression. Clin Exp Immunol. 2023;211: 176–183. doi:10.1093/cei/uxac123

18. Arpaia N, Green JA, Moltedo B, Arvey A, Hemmers S, Yuan S, et al. A Distinct Function of Regulatory T Cells in Tissue Protection. Cell. 2015;162: 1078–1089. doi:10.1016/j.cell.2015.08.021

19. Varanasi SK, Rajasagi N, Jaggi U, Rouse B. Role of IL-18 induced Amphiregulin expression on virus induced ocular lesions. Mucosal Immunol. 2018;11: 1705–1715. doi:10.1038/s41385-018-0058-8

20. Popovic B, Golemac M, Podlech J, Zeleznjak J, Bilic-Zulle L, Lukic ML, et al. IL-33/ST2 pathway drives regulatory T cell dependent suppression of liver damage upon cytomegalovirus infection. PLoS Pathog. 2017;13: e1006345. doi:10.1371/journal.ppat.1006345

21. Bai Y, Guan F, Zhu F, Jiang C, Xu X, Zheng F, et al. IL-33/ST2 Axis Deficiency Exacerbates Hepatic Pathology by Regulating Treg and Th17 Cells in Murine Schistosomiasis Japonica. J Inflamm Res. 2021;14: 5981–5998. doi:10.2147/JIR.S336404

22. Tariq M, Gallien S, Surenaud M, Wiedemann A, Jean-Louis F, Lacabaratz C, et al. Profound Defect of Amphiregulin Secretion by Regulatory T Cells in the Gut of HIV-Treated Patients. J Immunol. 2022;208: 2300–2308. doi:10.4049/jimmunol.2100725

23. Besnard A-G, Guabiraba R, Niedbala W, Palomo J, Reverchon F, Shaw TN, et al. IL-33-mediated protection against experimental cerebral malaria is linked to induction of type 2 innate lymphoid cells, M2 macrophages and regulatory T cells. PLoS Pathog. 2015;11: e1004607. doi:10.1371/journal.ppat.1004607

24. Jin RM, Warunek J, Wohlfert EA. Therapeutic administration of IL-10 and amphiregulin alleviates chronic skeletal muscle inflammation and damage induced by infection. Immunohorizons. 2018;2: 142–154. doi:10.4049/immunohorizons.1800024

25. Matta BM, Lott JM, Mathews LR, Liu Q, Rosborough BR, Blazar BR, et al. IL-33 is an unconventional Alarmin that stimulates IL-2 secretion by dendritic cells to selectively expand IL-33R/ST2+ regulatory T cells. J Immunol. 2014;193: 4010–4020. doi:10.4049/jimmunol.1400481

26. World Health Organization. Chagas disease. 13 Apr 2022 [cited 14 Jun 2022]. Available: https://www.who.int/news-room/fact-sheets/detail/chagas-disease-(american-trypanosomiasis)

27. Abrahamsohn IA, Coffman RL. Trypanosoma cruzi: IL-10, TNF, IFN-gamma, and IL-12 regulate innate and acquired immunity to infection. Exp Parasitol. 1996;84: 231–244. doi:10.1006/expr.1996.0109

28. Brener Z, Gazzinelli RT. Immunological control of Trypanosoma cruzi infection and pathogenesis of Chagas’ disease. Int Arch Allergy Immunol. 1997;114: 103–110. doi:10.1159/000237653

29. Golden JM, Tarleton RL. Trypanosoma cruzi: cytokine effects on macrophage trypanocidal activity. Exp Parasitol. 1991;72: 391–402. doi:10.1016/0014-4894(91)90085-b

30. Padilla AM, Bustamante JM, Tarleton RL. CD8+ T cells in Trypanosoma cruzi infection. Curr Opin Immunol. 2009;21: 385–390. doi:10.1016/j.coi.2009.07.006

31. Tarleton RL. Depletion of CD8+ T cells increases susceptibility and reverses vaccine-induced immunity in mice infected with Trypanosoma cruzi. J Immunol. 1990;144: 717– 724.

32. Guarner J. Chagas disease as example of a reemerging parasite. Semin Diagn Pathol. 2019;36: 164–169. doi:10.1053/j.semdp.2019.04.008

33. Lewis MD, Kelly JM. Putting Infection Dynamics at the Heart of Chagas Disease. Trends Parasitol. 2016;32: 899–911. doi:10.1016/j.pt.2016.08.009

34. de Carvalho JF, Lerner A. Fibromyalgia associated with Chagas’ disease treated with nutraceuticals. Clin Nutr ESPEN. 2021;42: 212–214. doi:10.1016/j.clnesp.2021.01.037

35. Köberle F. Chagas’ disease and Chagas’ syndromes: the pathology of American trypanosomiasis. Adv Parasitol. 1968;6: 63–116. doi:10.1016/s0065-308x(08)60472-8

36. Rincón-Acevedo CY, Parada-García AS, Olivera MJ, Torres-Torres F, Zuleta-Dueñas LP, Hernández C, et al. Clinical and Epidemiological Characterization of Acute Chagas Disease in Casanare, Eastern Colombia, 2012-2020. Front Med (Lausanne). 2021;8: 681635. doi:10.3389/fmed.2021.681635

37. Cossermelli W, Friedman H, Pastor EH, Nobre MR, Manzione A, Camargo ME, et al. Polymyositis in Chagas’s disease. Ann Rheum Dis. 1978;37: 277–280. doi:10.1136/ard.37.3.277

38. Laguens RP, Cossio PM, Diez C, Segal A, Vasquez C, Kreutzer E, et al. Immunopathologic and morphologic studies of skeletal muscle in Chagas’ disease. Am J Pathol. 1975;80: 153–162.

39. Buckner FS, Wilson AJ, Van Voorhis WC. Detection of live Trypanosoma cruzi in tissues of infected mice by using histochemical stain for beta-galactosidase. Infect Immun. 1999;67: 403–409. doi:10.1128/IAI.67.1.403-409.1999

40. Mendonça AAS, Gonçalves-Santos E, Souza-Silva TG, González-Lozano KJ, Caldas IS, Gonçalves RV, et al. Thioridazine aggravates skeletal myositis, systemic and liver inflammation in Trypanosoma cruzi-infected and benznidazole-treated mice. Int Immunopharmacol. 2020;85: 106611. doi:10.1016/j.intimp.2020.106611

41. Molina HA, Cardoni RL, Rimoldi MT. The neuromuscular pathology of experimental Chagas’ disease. J Neurol Sci. 1987;81: 287–300. doi:10.1016/0022-510x(87)90104-3

42. Monteón VM, Furuzawa-Carballeda J, Alejandre-Aguilar R, Aranda-Fraustro A, Rosales-Encina JL, Reyes PA. American trypanosomosis: in situ and generalized features of parasitism and inflammation kinetics in a murine model. Exp Parasitol. 1996;83: 267–274. doi:10.1006/expr.1996.0074

43. Weaver JD, Hoffman VJ, Roffe E, Murphy PM. Low-Level Parasite Persistence Drives Vasculitis and Myositis in Skeletal Muscle of Mice Chronically Infected with Trypanosoma cruzi. Infect Immun. 2019;87: e00081–19. doi:10.1128/IAI.00081-19

44. Araujo Furlan CL, Tosello Boari J, Rodriguez C, Canale FP, Fiocca Vernengo F, Boccardo S, et al. Limited Foxp3+ Regulatory T Cells Response During Acute Trypanosoma cruzi Infection Is Required to Allow the Emergence of Robust Parasite-Specific CD8+ T Cell Immunity. Front Immunol. 2018;9: 2555. doi:10.3389/fimmu.2018.02555

45. Tosello Boari J, Amezcua Vesely MC, Bermejo DA, Ramello MC, Montes CL, Cejas H, et al. IL-17RA signaling reduces inflammation and mortality during Trypanosoma cruzi infection by recruiting suppressive IL-10-producing neutrophils. PLoS Pathog. 2012;8: e1002658. doi:10.1371/journal.ppat.1002658

46. Ramirez-Archila MV, Muñiz J, Virgen-Ortiz A, Newton-Sánchez O, Melnikov VG, Dobrovinskaya OR. Trypanosoma cruzi: correlation of muscle lesions with contractile properties in the acute phase of experimental infection in mice (Mus musculus). Exp Parasitol. 2011;128: 301–308. doi:10.1016/j.exppara.2011.02.012

47. Tosello Boari J, Araujo Furlan CL, Fiocca Vernengo F, Rodriguez C, Ramello MC, Amezcua Vesely MC, et al. IL-17RA-Signaling Modulates CD8+ T Cell Survival and Exhaustion During Trypanosoma cruzi Infection. Front Immunol. 2018;9: 2347. doi:10.3389/fimmu.2018.02347

48. Koh B, Ulrich BJ, Nelson AS, Panangipalli G, Kharwadkar R, Wu W, et al. Bcl6 and Blimp1 reciprocally regulate ST2+ Treg-cell development in the context of allergic airway inflammation. J Allergy Clin Immunol. 2020;146: 1121–1136.e9. doi:10.1016/j.jaci.2020.03.002

49. Molofsky AB, Van Gool F, Liang H-E, Van Dyken SJ, Nussbaum JC, Lee J, et al. Interleukin-33 and Interferon-γ Counter-Regulate Group 2 Innate Lymphoid Cell Activation during Immune Perturbation. Immunity. 2015;43: 161–174. doi:10.1016/j.immuni.2015.05.019

50. Siede J, Fröhlich A, Datsi A, Hegazy AN, Varga DV, Holecska V, et al. IL-33 Receptor-Expressing Regulatory T Cells Are Highly Activated, Th2 Biased and Suppress CD4 T Cell Proliferation through IL-10 and TGFβ Release. PLoS One. 2016;11: e0161507. doi:10.1371/journal.pone.0161507

51. Vasanthakumar A, Chisanga D, Blume J, Gloury R, Britt K, Henstridge DC, et al. Sex-specific adipose tissue imprinting of regulatory T cells. Nature. 2020;579: 581–585. doi:10.1038/s41586-020-2040-3

52. Vasanthakumar A, Moro K, Xin A, Liao Y, Gloury R, Kawamoto S, et al. The transcriptional regulators IRF4, BATF and IL-33 orchestrate development and maintenance of adipose tissue-resident regulatory T cells. Nat Immunol. 2015;16: 276–285. doi:10.1038/ni.3085

53. Muñoz-Rojas AR, Mathis D. Tissue regulatory T cells: regulatory chameleons. Nat Rev Immunol. 2021;21: 597–611. doi:10.1038/s41577-021-00519-w

54. Abbas AK, Trotta E, R Simeonov D, Marson A, Bluestone JA. Revisiting IL-2: Biology and therapeutic prospects. Sci Immunol. 2018;3: eaat1482. doi:10.1126/sciimmunol.aat1482

55. Griesenauer B, Paczesny S. The ST2/IL-33 Axis in Immune Cells during Inflammatory Diseases. Front Immunol. 2017;8: 475. doi:10.3389/fimmu.2017.00475

56. de Oliveira MFA, Talvani A, Rocha-Vieira E. IL-33 in obesity: where do we go from here? Inflamm Res. 2019;68: 185–194. doi:10.1007/s00011-019-01214-2

57. Li C, Wang G, Sivasami P, Ramirez RN, Zhang Y, Benoist C, et al. Interferon-α-producing plasmacytoid dendritic cells drive the loss of adipose tissue regulatory T cells during obesity. Cell Metab. 2021;33: 1610–1623.e5. doi:10.1016/j.cmet.2021.06.007

58. Morrow KN, Coopersmith CM, Ford ML. IL-17, IL-27, and IL-33: A Novel Axis Linked to Immunological Dysfunction During Sepsis. Front Immunol. 2019;10: 1982. doi:10.3389/fimmu.2019.01982

59. Junqueira C, Caetano B, Bartholomeu DC, Melo MB, Ropert C, Rodrigues MM, et al. The endless race between Trypanosoma cruzi and host immunity: lessons for and beyond Chagas disease. Expert Rev Mol Med. 2010;12: e29. doi:10.1017/S1462399410001560

60. Paroli AF, Gonzalez PV, Díaz-Luján C, Onofrio LI, Arocena A, Cano RC, et al. NLRP3 Inflammasome and Caspase-1/11 Pathway Orchestrate Different Outcomes in the Host Protection Against Trypanosoma cruzi Acute Infection. Front Immunol. 2018;9: 913. doi:10.3389/fimmu.2018.00913

61. Endo Y, Karvar M, Sinha I. Muscle Cryoinjury and Quantification of Regenerating Myofibers in Mice. Bio Protoc. 2021;11: e4036. doi:10.21769/BioProtoc.4036

62. Klose CSN, Artis D. Innate lymphoid cells as regulators of immunity, inflammation and tissue homeostasis. Nat Immunol. 2016;17: 765–774. doi:10.1038/ni.3489

63. Tait Wojno ED, Beamer CA. Isolation and Identification of Innate Lymphoid Cells (ILCs) for Immunotoxicity Testing. Methods Mol Biol. 2018;1803: 353–370. doi:10.1007/978-1-4939-8549-4_21

64. Molofsky AB, Savage AK, Locksley RM. Interleukin-33 in Tissue Homeostasis, Injury, and Inflammation. Immunity. 2015;42: 1005–1019. doi:10.1016/j.immuni.2015.06.006

65. Verri WA, Guerrero ATG, Fukada SY, Valerio DA, Cunha TM, Xu D, et al. IL-33 mediates antigen-induced cutaneous and articular hypernociception in mice. Proc Natl Acad Sci U S A. 2008;105: 2723–2728. doi:10.1073/pnas.0712116105

66. Tarleton RL. CD8+ T cells in Trypanosoma cruzi infection. Semin Immunopathol. 2015;37: 233–238. doi:10.1007/s00281-015-0481-9

67. Chen X, Li M, Chen B, Wang W, Zhang L, Ji Y, et al. Transcriptome sequencing and analysis reveals the molecular mechanism of skeletal muscle atrophy induced by denervation. Ann Transl Med. 2021;9: 697. doi:10.21037/atm-21-1230

68. Sastourné-Arrey Q, Mathieu M, Contreras X, Monferran S, Bourlier V, Gil-Ortega M, et al. Adipose tissue is a source of regenerative cells that augment the repair of skeletal muscle after injury. Nat Commun. 2023;14: 80. doi:10.1038/s41467-022-35524-7

69. Yan Z, Choi S, Liu X, Zhang M, Schageman JJ, Lee SY, et al. Highly coordinated gene regulation in mouse skeletal muscle regeneration. J Biol Chem. 2003;278: 8826–8836. doi:10.1074/jbc.M209879200

70. Cayrol C, Girard J-P. Interleukin-33 (IL-33): A critical review of its biology and the mechanisms involved in its release as a potent extracellular cytokine. Cytokine. 2022;156: 155891. doi:10.1016/j.cyto.2022.155891

71. Liew FY, Girard J-P, Turnquist HR. Interleukin-33 in health and disease. Nat Rev Immunol. 2016;16: 676–689. doi:10.1038/nri.2016.95

72. Lüthi AU, Cullen SP, McNeela EA, Duriez PJ, Afonina IS, Sheridan C, et al. Suppression of interleukin-33 bioactivity through proteolysis by apoptotic caspases. Immunity. 2009;31: 84–98. doi:10.1016/j.immuni.2009.05.007

73. Cohen ES, Scott IC, Majithiya JB, Rapley L, Kemp BP, England E, et al. Oxidation of the alarmin IL-33 regulates ST2-dependent inflammation. Nat Commun. 2015;6: 8327. doi:10.1038/ncomms9327

74. Braband KL, Kaufmann T, Floess S, Zou M, Huehn J, Delacher M. Stepwise acquisition of unique epigenetic signatures during differentiation of tissue Treg cells. Front Immunol. 2022;13: 1082055. doi:10.3389/fimmu.2022.1082055

75. Frisbee AL, Saleh MM, Young MK, Leslie JL, Simpson ME, Abhyankar MM, et al. IL-33 drives group 2 innate lymphoid cell-mediated protection during Clostridium difficile infection. Nat Commun. 2019;10: 2712. doi:10.1038/s41467-019-10733-9

76. Ngo Thi Phuong N, Palmieri V, Adamczyk A, Klopfleisch R, Langhorst J, Hansen W, et al. IL-33 Drives Expansion of Type 2 Innate Lymphoid Cells and Regulatory T Cells and Protects Mice From Severe, Acute Colitis. Front Immunol. 2021;12: 669787. doi:10.3389/fimmu.2021.669787

77. Uddin MJ, Leslie JL, Burgess SL, Oakland N, Thompson B, Abhyankar M, et al. The IL-33-ILC2 pathway protects from amebic colitis. Mucosal Immunol. 2022;15: 165–175. doi:10.1038/s41385-021-00442-2

78. Aparicio-Domingo P, Cannelle H, Buechler MB, Nguyen S, Kallert SM, Favre S, et al. Fibroblast-derived IL-33 is dispensable for lymph node homeostasis but critical for CD8 T-cell responses to acute and chronic viral infection. Eur J Immunol. 2021;51: 76–90. doi:10.1002/eji.201948413

79. Bonilla WV, Fröhlich A, Senn K, Kallert S, Fernandez M, Johnson S, et al. The alarmin interleukin-33 drives protective antiviral CD8^+^ T cell responses. Science. 2012;335: 984– 989. doi:10.1126/science.1215418

80. Marx A-F, Kallert SM, Brunner TM, Villegas JA, Geier F, Fixemer J, et al. The alarmin interleukin-33 promotes the expansion and preserves the stemness of Tcf-1+ CD8+ T cells in chronic viral infection. Immunity. 2023;56: 813–828.e10. doi:10.1016/j.immuni.2023.01.029

81. Belkaid Y, Tarbell K. Regulatory T cells in the control of host-microorganism interactions (*). Annu Rev Immunol. 2009;27: 551–589. doi:10.1146/annurev.immunol.021908.132723

82. Rocha IH, Ferreira Marques AL, Moraes GV, Alves da Silva DA, Silva MV da, Rodrigues V, et al. Metabolic and immunological evaluation of patients with indeterminate and cardiac forms of Chagas disease. Medicine (Baltimore). 2020;99: e23773. doi:10.1097/MD.0000000000023773

83. Albareda MC, Olivera GC, Laucella SA, Alvarez MG, Fernandez ER, Lococo B, et al. Chronic human infection with Trypanosoma cruzi drives CD4+ T cells to immune senescence. J Immunol. 2009;183: 4103–4108. doi:10.4049/jimmunol.0900852

84. Poncini CV, Giménez G, Pontillo CA, Alba-Soto CD, de Isola ELD, Piazzón I, et al. Central role of extracellular signal-regulated kinase and Toll-like receptor 4 in IL-10 production in regulatory dendritic cells induced by Trypanosoma cruzi. Mol Immunol. 2010;47: 1981– 1988. doi:10.1016/j.molimm.2010.04.016

85. Piron M, Fisa R, Casamitjana N, López-Chejade P, Puig L, Vergés M, et al. Development of a real-time PCR assay for Trypanosoma cruzi detection in blood samples. Acta Trop. 2007;103: 195–200. doi:10.1016/j.actatropica.2007.05.019

86. Fiocca Vernengo F, Beccaria CG, Araujo Furlan CL, Tosello Boari J, Almada L, Gorosito Serrán M, et al. CD8+ T Cell Immunity Is Compromised by Anti-CD20 Treatment and Rescued by Interleukin-17A. mBio. 2020;11: e00447–20. doi:10.1128/mBio.00447-20

87. Souza DG, Cara DC, Cassali GD, Coutinho SF, Silveira MR, Andrade SP, et al. Effects of the PAF receptor antagonist UK74505 on local and remote reperfusion injuries following ischaemia of the superior mesenteric artery in the rat. Br J Pharmacol. 2000;131: 1800– 1808. doi:10.1038/sj.bjp.0703756

88. Frankish A, Diekhans M, Ferreira A-M, Johnson R, Jungreis I, Loveland J, et al. GENCODE reference annotation for the human and mouse genomes. Nucleic Acids Res. 2019;47: D766–D773. doi:10.1093/nar/gky955

89. Patro R, Duggal G, Love MI, Irizarry RA, Kingsford C. Salmon provides fast and bias-aware quantification of transcript expression. Nat Methods. 2017;14: 417–419. doi:10.1038/nmeth.4197

90. Chen EY, Tan CM, Kou Y, Duan Q, Wang Z, Meirelles GV, et al. Enrichr: interactive and collaborative HTML5 gene list enrichment analysis tool. BMC Bioinformatics. 2013;14: 128. doi:10.1186/1471-2105-14-128

91. Kuleshov MV, Jones MR, Rouillard AD, Fernandez NF, Duan Q, Wang Z, et al. Enrichr: a comprehensive gene set enrichment analysis web server 2016 update. Nucleic Acids Res. 2016;44: W90–97. doi:10.1093/nar/gkw377

92. Xie Z, Bailey A, Kuleshov MV, Clarke DJB, Evangelista JE, Jenkins SL, et al. Gene Set Knowledge Discovery with Enrichr. Current Protocols. 2021;1: e90. doi:10.1002/cpz1.90

93. Deacon RMJ. Measuring the strength of mice. J Vis Exp. 2013; 2610. doi:10.3791/2610

